# Diagnostic Evidence GAuge of Single cells (DEGAS): A flexible deep-transfer learning framework for prioritizing cells in relation to disease

**DOI:** 10.1101/2020.06.16.142984

**Authors:** Travis S. Johnson, Christina Y. Yu, Zhi Huang, Siwen Xu, Tongxin Wang, Chuanpeng Dong, Wei Shao, Mohammad Abu Zaid, Xiaoqing Huang, Yijie Wang, Christopher Bartlett, Yan Zhang, Brian A. Walker, Yunlong Liu, Kun Huang, Jie Zhang

## Abstract

We propose *DEGAS* (Diagnostic Evidence GAuge of Single cells), a novel deep transfer learning framework, to transfer disease information from patients to cells. We call such transferrable information “impressions,” which allow individual cells to be associated with disease attributes like diagnosis, prognosis, and response to therapy. Using simulated data and ten diverse single cell and patient bulk tissue transcriptomic datasets from Glioblastoma Multiforme (GBM), Alzheimer’s Disease (AD), and Multiple Myeloma (MM), we demonstrate the feasibility, flexibility, and broad applications of the *DEGAS* framework. *DEGAS* analysis on newly generated myeloma single cell transcriptomics led to the identification of *PHF19^high^* myeloma cells associated with progression.

## Background

The emergence of single cell RNA sequencing (scRNA-seq) in 2009 has revolutionized the medical research community with single cell level resolution, providing a much deeper understanding of transcriptomic heterogeneity in tissues and diseases. Now that scRNA-seq is a standard part of the biomedical research toolbox, increasing numbers of scRNA-seq studies have been published [1, 2], and databases have quickly accumulated with scRNA-seq data, such as Hemberg lab [3], scRNASeqDB [4], SCPortalen [5], Allen Institute Cell Types Database, and the NCBI Gene Expression Omnibus (GEO) [5]. Many methods have been developed to analyze scRNA-seq data, the most notable being *Seurat*, which includes ways to cluster and normalize cell expression as well as perform integrative analysis with other data types (*e.g.*, CITE-seq and ATAC-seq) [6]. These methods are important for understanding many prognostic and diagnostic disease attributes in scRNA-seq data. Here we use “disease attributes” as a broad term inclusive of many types of information and labeling such as diagnostic information, disease subtypes, disease status, prognostic information like survival, and responses to therapy. For *Seurat* and similar methods, while cell types/clusters can be identified and associated with disease attributes [7–10], individual cells are unable to be associated in the same manner. This may result in failing to identify subsets of cells associated with disease attributes, especially if the disease-associated cells cluster together with non-disease-associated cells.

Currently, disease associated cell types can be identified by transferring molecular heterogeneity information from cells to patients using single cell expression deconvolution [11–13]. However, this approach is limited as it focuses on the changes in relative abundance of subtypes of cells instead of transcription changes of these cells. The resolution of the cell subtyping is constrained by the clustering experiment. Therefore, novel machine learning methods that can transfer information from patients to cells and identify latent links between them are sorely needed to leverage the relative strengths of single cell and patient level data. For example, in cancer studies, bulk transcriptomic data is ideal for studying inter-tumor heterogeneity and scRNA-seq is ideal for studying intra-tumor heterogeneity. However, such integration faces numerous challenges since different data modalities and different data sources can have different characteristics in terms of quantity, quality, distribution and resolution [1]. For instance, it is common to find studies with a large number of patient samples for bulk tissue RNA sequencing (RNA-seq), whereas studies with scRNA-seq data usually contain a small number of patient samples. Most scRNA-seq experiments generate a large number of cells per sample, making the scaling of such data to multiple tissue samples computationally difficult [1]. On the other hand, a large patient sample size is often required for statistical studies such as prediction of disease attributes [14]. If traditional methods were used, the resulting scRNA-seq data could end up with cell numbers on the scale of millions making such studies more difficult.

To address these challenges, previous studies have directly established associations of diseases with cell types derived from scRNA-seq without using deconvolution. These methods mainly utilize unsupervised methods and focused primarily on the number of differentially expressed genes (DEGs) in a given cell type corresponding to DEGs related to some disease attribute [15, 16]. For example, Gawel *et al.* used enrichment of the cell cluster specific DEGs and multicellular disease models (MCDMs) to visualize the cell types for prioritization [7]. *Muscat* identified DEGs between treatment groups in scRNA-seq samples which were used to identify cell types related to sample treatment [17]. Alternatively, k nearest neighbor (*kNN*) graphs have been used to identify cell types that undergo transcriptional changes related to biological perturbations [18]. The cell type prioritization tool *Augur* did not primarily rely on DEGs, but still focused the biological resolution to the cell type level [19]. They trained classifiers on each cell type with respect to the disease state of the tissue from which those cells were sampled. The accuracy of the classifier in each cell type was used to prioritize its relation to the disease state of interest [19]. These methods rely on either prior knowledge to calculate enrichment of DEGs or require scRNA-seq data from both disease and normal samples. Furthermore, all of these existing methods are reliant on accurately defining the cell types within a scRNA-seq experiment. In summary, these methods assign disease associations to the previously defined cell types and not to the individual cells.

To address such challenges as prioritizing individual cells in relation to disease with considerations on sample size and computational cost, we established the combined deep learning and transfer learning framework called *DEGAS* (<u>D</u>iagnostic <u>E</u>vidence <u>GA</u>uge of <u>S</u>ingle cells) to integrate scRNA-seq and bulk tissue transcriptomic data with the goal to transfer clinical information from patients to cells. The ability of *DEGAS* to assign patient-level disease attributes to single cells, among other functions, provides a flexible and useful tool to prioritize cells, cell types, patients, and patient subtypes in relation to disease attributes. In this paper, we focus on the most relevant use case of associating disease attributes from patients to individual cells since there is no current state-of-the art technique to perform this task.

We use transcriptomic data as an example where bulk expression is referred to as patients and scRNA-seq is referred to as cells. The rationale behind the *DEGAS* framework is that scRNA-seq data and patient-level transcriptomic data (*e.g.*, RNA-seq with clinical information) share the same feature space (*i.e.*, common set of genes). In addition, a natural connection exists between the two data types that can be leveraged to further identify the associations between patients and cells. Viewing this association as a graph (Fig. 1), we can connect the disease attributes in patients to individual cells, via a latent representation of the common feature space (selected genes). This latent representation fitting two datasets can be learned using a transfer learning technique called domain adaptation [20–23]. Domain adaptation applies linear or non-linear transformations on the features for both datasets so that their distributions are similar after the transformations. Our biological intuition is thus: the expression patterns of genes in cells and tissues should carry a portion of the same biological patterns such as molecular pathways, signaling cascades, and/or metabolic processes, making the information learned from this portion of gene expression patterns transferable between patients and cells. Our hypothesis is that the latent representation learned from these shared gene expression patterns will be simultaneously predictive of patient disease attributes and cellular subtypes. Similar hypotheses are already adopted to transfer information between different single cell experiments [6, 24–28] and to transfer information from bulk transcriptomic cell type atlases to single cell experiments [29].

**Fig. 1.**
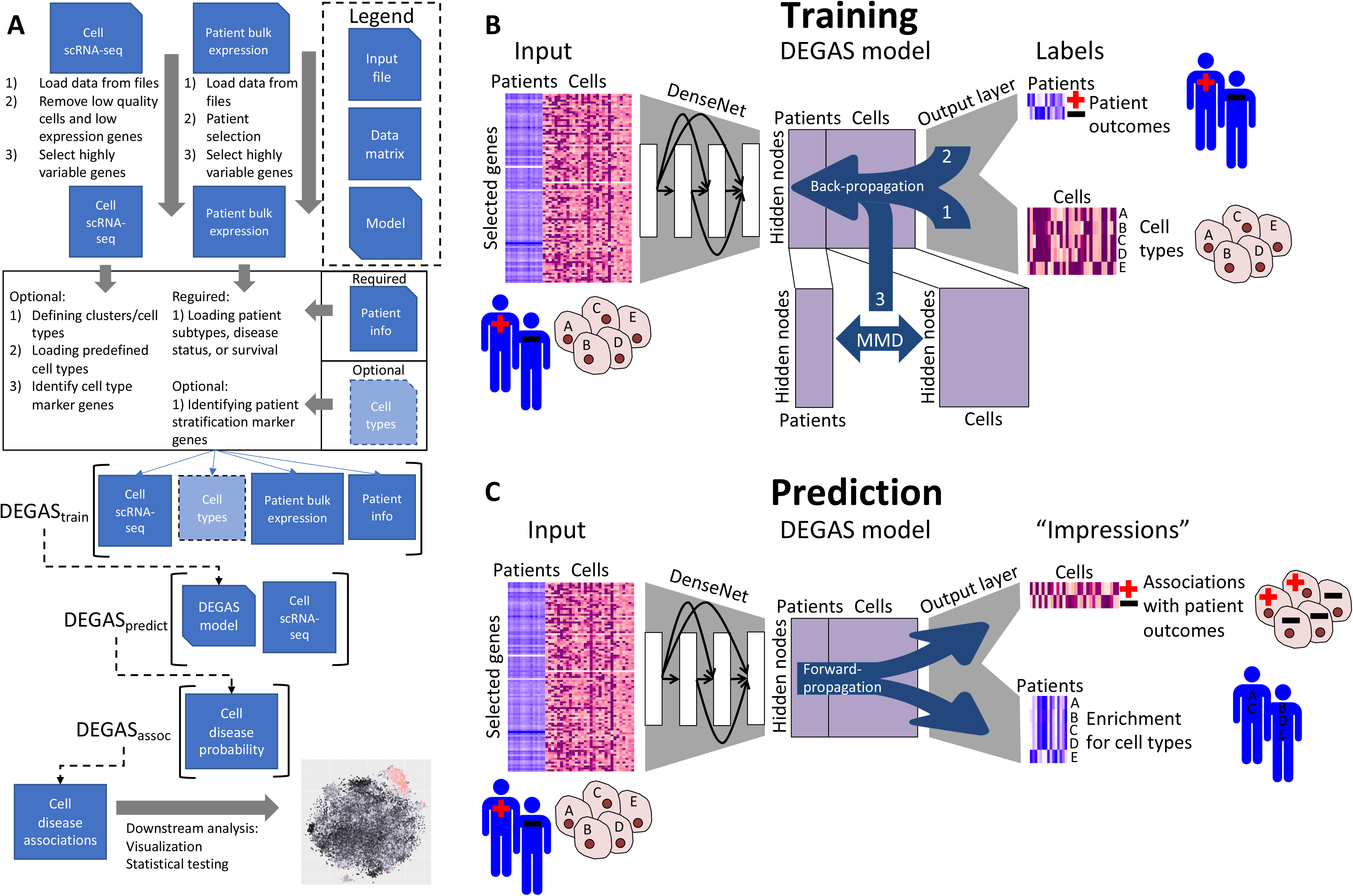
A workflow diagram of the *DEGAS* framework. **A**) The workflow for a typical experiment with *DEGAS*. Note that *DEGAS* is not meant to replace the abundant packages available to load, preprocess, select features, cluster, and visualize scRNA-seq data. It is rather meant to augment these packages to assign disease associations to cells. **B**) The scRNA-seq and patient expression data are preprocessed into expression matrices. Next, Bootstrap aggregated DenseNet *DEGAS* models are trained using both single cell and patient disease attributes using a multitask learning neural network that learns latent representation reducing the differences between patients and single cells at the final hidden layer using maximum mean discrepancy (MMD). **C**) The output layer of this model can be used to simultaneously infer disease attribute impressions in single cells and cellular composition impressions in patients.

In our *DEGAS* framework, we incorporate patient-level disease attributes information with cell type information from disparate datasets to perform cell prioritization on scRNA-seq data. These disease associations in cells can be attributed to disease-related biological perturbations identified in the patients. This novel deep transfer learning approach simultaneously trains a model on single cell data and patient data along with their labels and learns a representation in which the cells and patients occupy the same latent space. Multitask learning, also known as parallel transfer learning, is precisely designed to achieve these two goals. Used extensively in computer vision, multitask learning learns a low dimensional representation of the input data to optimally address multiple tasks. Examples of such application in medical science include predicting benign versus malignant tumor samples and subclassification in breast cancer histology images [30, 31]. In this paper, we further extend this line of research to include datasets with patient disease attributes that can be trained simultaneously so that the disease attributes can be transferred (or cross-mapped) between single cells and patients. Specifically, our framework enables knowledge learned from patients using deep learning models to be transferred to single cells and vice versa. The major advantages of our transfer learning framework are that the single-cell gene expression data and clinical bulk gene expression data can come from different patient cohorts of the same disease without matched data while the disease associations can still be directly assigned to individual cells. This flexibility not only presents an ingenious way to integrate molecular omics data analysis in different levels, but also virtually merges them into the same cohort, which makes studying a broad variety of heterogeneous diseases possible.

Various types of workflows can integrate the *DEGAS* framework, which can be tailored to user preference and data availability. These workflows consist of preprocessing, formatting data, training *DEGAS* models using the *DEGAS* framework, predicting disease associations in cells using the *DEGAS* framework, and downstream analysis (Fig. 1A). The *DEGAS* framework in its simplest form can be broken into three tasks during model training: 1) correctly labeling cells with a cellular subtype using multitask learning; 2) correctly assigning clinical labels to patients using multitask learning; and 3) generating a latent space in which patients and cells are comparable using domain adaptation (Fig. 1B). To perform *DEGAS* analysis, first we select representative gene features that are predictive of cell type, predictive of patient disease attributes, and present at measurable levels in both scRNA-seq and bulk transcriptomic data. Secondly, we apply deep learning models to learn the latent representation of the single-cell and patient-level transcriptomic data, with the goal to simultaneously minimize cell type classification error, patient disease attribute prediction error, and the differences between cells and patients in their latent representation. Finally, the patient-level disease attributes such as survival and clinical subtypes is predicted in the single cells using the patient label output layer and cell types are predicted in patients using the cell type output layer (Fig. 1C). We call these transferrable label probabilities “impressions” since information from gene expression of disparate data types and studies can be extracted and the characteristics from one data type can be mapped to another. These impressions of disease attributes in single cells can be wide ranging characteristics of the patient samples but must be categorical or time to event. The most interesting of them that can be used in *DEGAS* are disease status, disease subtype, survival, and response to therapy. Disease status, subtype, and survival were used in our current experiments but there would also be much utility in identifying cells associated with poor response to treatment as the data become available. Furthermore, we emphasize the ability to make predictions of patient disease attributes in individual cells since there is a lack of such method to perform this task to the best of our knowledge. *DEGAS* is developed as a generalizable model generating deep transfer learning framework that can be applied to any disease data as long as the data contain clinical information for a cohort of patients or a separate clustering analysis result on sets of cells from single cell level omic experiments of the same disease. Since there is not an inherent limitation to the use of transcriptomic data, *DEGAS* can be potentially expanded to accommodate other modalities of data with proper normalization steps.

To demonstrate the feasibility and effectiveness of the *DEGAS* framework, we first tested it on simulated data and glioblastoma (GBM) transcriptomic data, which contain ground-truth labels of cell types on single cell gene expression data and clinical labels for patient bulk tissue gene expression data. Then we applied *DEGAS* to multiple Alzheimer’s disease (AD) gene expression datasets from Mount Sinai/JJ Peters VA Medical Center, Allen Institute, Grubmann *et al.* [32], and Mathys et al. [15] in which certain cell type changes (microglia and neuron) are largely known [33–39]. Finally, as an exploratory tool, we applied *DEGAS* to study multiple myeloma (MM) transcriptomic data, where the disease associated subtypes of cells are largely unknown.

MM is a late stage of myeloma that stems from the proliferation of aberrant clonal plasma cells in the bone marrow that secrete monoclonal immunoglobulins and is the second most common blood cancer in the United States [40]. Patient level transcriptomic data for MM has been widely available for some time and has been used to identify subtypes of MM with different prognoses [41]. However, only recently has scRNA-seq become available for MM [9, 42, 43] and few studies have identified the most high-risk subtypes of cells [9]. Here we combined our newly generated late-stage myeloma scRNA-seq data from four local samples and bulk tissue data from the Multiple Myeloma Research Foundation CoMMpass study, and then applied *DEGAS* to infer clinical impressions for myeloma cell subtypes and successfully identified a *PHF19^high^* myeloma cell subgroup associated with a high-risk of progression.

## Methods

### Experimental design and datasets

For a *DEGAS* cell prioritization experiment, one scRNA-seq dataset, one bulk expression dataset, and patient sample labels (matched with the bulk data samples) are required as input. After feature selection and scaling (see ***Feature selection and scaling***) of the raw input expression data, there should be two expression matrices with rows corresponding to samples/cells and matching columns corresponding to genes. The bulk patient sample labels should be one-hot encoded in a matrix with rows corresponding to each sample and the columns corresponding to each class of label. For survival sample labels the first column should be time and the second column should be the even indicator (1 event and 0 censored). If cell labels are also available, they should also be one-hot encoded with each row corresponding to a cell and each column corresponding to a class of label. The *DEGAS* models can be trained and predicted on these formatted data (Fig. 1A).

In this study we analyzed simulated data and data from three different diseases, GBM, AD, and MM, to test the *DEGAS* framework and apply it for novel discoveries. The simulation, GBM, and AD experiments were primarily used as validation datasets since the ground truth is known. The simulated data were generated so that cell types are directly related to disease status in patients. For GBM data, we used scRNA-seq data for five tumors from Patel *et al.* [44] and microarray data for the GBM TCGA cohort [45] (Table 1). For AD data, we used human scRNA-seq from Allen Institute for Brain Science (AIBS) Cell Types Database (https://celltypes.brain-map.org/) and AD patient RNA-seq from the Mount Sinai/JJ Peters VA Medical Center Brain Bank (MSBB) study[46] (Table 1).

**Table 1.** Summary of the clinical features in each patient cohorts used in training. * Final age category is >90 years.

| Glioblastoma Multiforme TCGA |  |
| --- | --- |
| Feature | Details |
| Sex | 74 Male, 37 Female |
| Age (years) | Range: 14-83, Mean: 56, Median: 58 |
| Clinical GBM subtype | 34 Classical, 33 Mesenchymal, 9 Neural, 35 Proneural |
| Alzheimer's Disease MSBB |  |
| Feature | Details |
| Sex | 90 Male, 131 Female |
| Age (years) | Range: 61-90+, Mean* > 82, Median = 84 |
| AD diagnosis | 135 AD, 86 Control |
| **Multiple Myeloma MMRF** |  |
| **Feature** | **Details** |
| Sex | 387 Male, 260 Female |
| Age (years) | Range: 27-93, Mean: 64, Median: 64 |
| Relapse-free survival time (days) | Range: 13-1753, Mean: 665.4, Median: 629<br>200 patients progressed |

We further expanded our inquiry into MM, which served as a discovery study. Since the plasma cell subtypes are less understood in relation to MM clinical outcomes, we aimed to identify subtypes of plasma cells associated with worse prognosis. We first utilized 647 CD138^+^-enriched bone marrow patient samples from the Multiple Myeloma Research Foundation CoMMpass study (MMRF). These data were generated as part of the Multiple Myeloma Research Foundation Personalized Medicine Initiatives (https://research.themmrf.org). The dataset consisted of tumor tissue RNA-seq data and corresponding clinical information including progression free survival (PFS) time and survival status. PFS was defined as the time taken for a patient to relapse, progress, or die after treatment of the initial tumor. The demographic information of the MMRF patients are shown in Table 1. The first scRNA-seq data used in this study were generated by us using samples consisting of CD138^+^ plasma cells purified from bone marrow from four myeloma patients including two MM patients.

There were six total samples collected from myeloma patients. Of these, four samples passed initial quality control checks. Sample 1 and 6 were dropped due to sample degradation and data quality issues. This in turn left with four usable samples, *i.e.*, samples 2, 3, 4, and 5 for our study. The low number of patients was a good test case considering most scRNA-seq experiments frequently have few patients. The single cells were sequenced using 10x Genomics and Illumina NovaSeq6000 sequencer. *CellRanger 2.1.0* (http://support.10xgenomics.com/) was utilized to process the raw sequence data. Briefly, *CellRanger* used *bcl2fastq* (https://support.illumina.com/) to demultiplex raw base sequence calls generated from the sequencer into sample-specific FASTQ files. The FASTQ files were then aligned to the human reference genome GRCh38 with RNA-seq aligner *STAR*. The aligned reads were traced back to individual cells and the gene expression level of individual genes were quantified based on the number of UMIs (unique molecular indices) detected in each cell. The filtered gene-cell barcode matrices generated by *CellRanger* were used for further analysis. A second publicly available myeloma scRNA-seq dataset was used for validation, which consisted of NHIP (normal control), MGUS (monoclonal gammopathy of undetermined significance), SMM (smoldering multiple myeloma), and MM [42]. A second bulk tissue dataset was used for validating the proportional hazards modeling. This dataset consisted of bulk expression profiling by microarray of CD138+ plasma cells with overall survival (OS) information for 559 MM patients [41]. The detailed information of the four datasets is shown in Table 2.

**Table 2.** Overview of all datasets used in the analysis. *The simulated patients were generated from the splatter simulated cells by combining known proportions of cell types. “None” is used to denote the lack of labels for the cells/samples in a given dataset. ^†^Cells were down-sampled from the total number of cells because some cell types were over-represented.

| Study | Dataset | Sample size | Data type | Attribute |
| --- | --- | --- | --- | --- |
| **Simulation** | Simulated cells* | 5000 cells | scRNA-seq | Cell type |
|  | Simulated patients* | 600 patients | RNA-seq | Disease status |
| **Glioblastoma** | Patel *et al.*, 2014 | 532 cells<br>(5 patients) | scRNA-seq<br>(SMART-seq) | None |
|  | TCGA GBM | 111 patients | Microarray | GBM subtype |
| **Alzheimer's disease** | AIBS | 47,396 cells<br>(11 patients) | scRNA-seq<br>(SMART-seq) | Brain cell types |
|  | Grubman *et al.*, 2019 | 13,214 cells<br>(12 patients) | snRNA-seq<br>(10x Genomics) | AD and normal brain cell types |
|  | Mathys *et al.*, 2019 | 5288 cells<sup>†</sup><br>(48 patients) | snRNA-seq<br>(10x Genomics) | AD and normal brain cell types |
|  | MSBB | 682 samples<br>(221 patients) | RNA-seq | AD diagnosis |
| **Multiple myeloma** | MMRF | 647 patients | RNA-seq | PFS |
|  | IUSM | 22,968 cells<br>(4 patients) | scRNA-seq<br>(10x Genomics) | Subtype cluster<br>(Subtype 1-5) |
|  | Ledergor *et al.*, 2019 | 13,440 cells<br>(35 patients) | scRNA-seq<br>(MARS-seq) | Malignancy<br>(NHIP, MGUS, SMM, MM) |
|  | Zhan *et al.*, 2006 | 559 patients | Microarray | OS |

### Transfer learning using DEGAS

Several types of labels including Cox proportional hazards, patient classification, and cell type classification, along with maximum mean discrepancy (MMD), a technique used to match distributions across different sets of data [22], were combined to create the multitask transfer learning framework *DEGAS*.

The first step was to find a set of gene expression features that were both informative of cell type and of patient disease attribute (*e.g.*, recurrence). The intersection of high variance genes found in the scRNA-seq and bulk expression data of patient samples are used for further analysis. The definition of this gene set is up to the user but *Seurat-CCA*, *LASSO* selection, and even statistical tests such as t-test and f-test can be used to define the gene set. Since these features are the same between patients and single cells, the patients and cells share the same input layer. This makes it possible to predict proportional hazard and cell type regardless of the input sample type (patient or single cell data).

All experiments in this manuscript use a five-bootstrap aggregated three-layer DenseNet-based implementation of *DEGAS*, but the simplest form of the *DEGAS* framework is a single layer network. In our description of the overall architecture below (shown in Fig. 1B,C), we used a single layer network for the purpose of simplicity. The following Eq. 1 can nevertheless be extrapolated to multiple layers and architectures, some of which we have already included in our open-source software package. First, a hidden layer was used to transform the genes into a lower dimension using a sigmoid activation function (Eq. 1). Where *X* represents an input expression matrix, *θ_Hidden_* represents the hidden layer weights, and *b_Hidden_* represents the hidden layer bias.

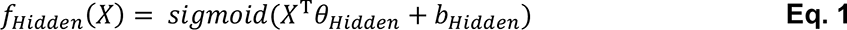

Next, output layers were added for both the patient output and for the single cell output. For the single cells, there could be classification output or no output. No output means there are no known labels for the single cells to match. Similarly, patients could have Cox proportional hazard output, classification output, or no output (implying no known labels for patients).

The Cox proportional hazards estimates consisted of a linear transformation to a single output followed by a sigmoid activation function (Eq. 2):

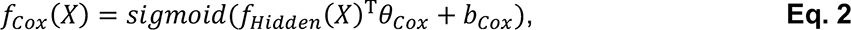

where the variable *X* represents an input expression matrix, *θ_Cox_* represents the Cox proportional hazard layer weights [47], and *b_Cox_* represents the Cox proportional hazard layer bias. The classification output consisted of a transformation to the same number of outputs as the number of labels, *i.e.*, patient subtypes, cellular subtypes, using a softmax activation function (Eq. 3).

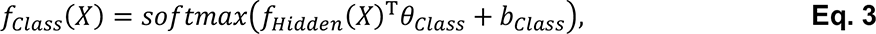

*θ_Class_* represents the classification layer weights and *b_Class_* represents the classification layer bias.

To train the *DEGAS* model, we need to compute three types of loss functions for the Cox proportional hazards output, classification output, and MMD [22] respectively. The Cox proportional hazards loss [47] was calculated only for the patient expression data (*X_Pat_*) using the followup period (*C*), and event status (*t*) (Eq. 4). Similarly, the patient classification loss was only calculated for the patient data (*X_Pat_*) using the patient labels (*Y_Pat_*). Alternatively, the cellular classification loss was only calculated for the single cell expression data (*X_Cell_*) and true subtype label (*Y_Cell_*) (Eq. 5). The MMD loss was calculated between the patient expression data (*X_Pat_*) and the single cell expression data (*X_Cell_*) (Eq. 6), which is the key for mapping the distributions of the data representations between the single-cell and patient bulk tissue data.

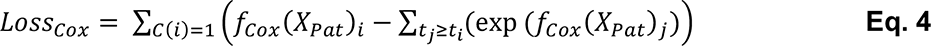

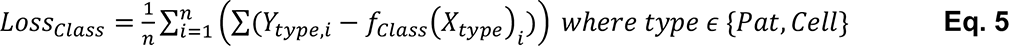

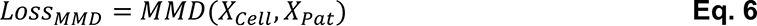

Besides the three losses, we also add a *L_2_*-regularization loss term to constrain for the complexity of the model. The overall loss function was the weighted sum of the four types of loss using the hyper-parameters *λ*_0_ (single cell loss function), *λ*_1_ (patient loss function), *λ*_2_ (MMD loss), and *λ*_3_ (regularization loss), so that the importance of each loss term and regularization term could be adjusted (Eq. 7):

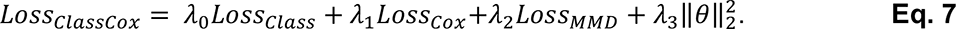

To address more diverse scenarios, we can also adapt Eq. 7 for two classification outputs (Eq. 8), a single classification output without patient disease attribute (Eq. 9), a single classification output without cell type label (Eq. 10), or a single Cox output without cell type label (Eq. 11):

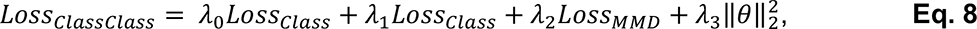

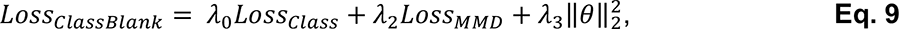

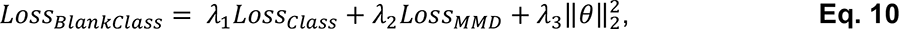

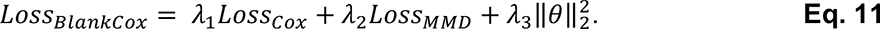

In summary, a common hidden layer was used to merge the single cells and patient data. Next, an output layer was added to predict the proportional hazards or classes of the patient samples [47]. The loss function for the proportional hazards prediction or patient classification was back-propagated across both layers for each patient. The single cells also had an output layer consisting of a softmax output to predict the cellular subtype of each cell. Error was back-propagated across both layers from the label output for each cell. Finally, a model was learned that can model both the single cells and the patients. To perform this task, we utilized the MMD method [22] to reduce the differences between patients and cells in a low dimensional representation. Both single cell and patient bulk tissue data were combined into a single group such that the MMD loss was minimized between patient bulk tissue data and single cell data from multiple patients. Because there are many different combinations of these outputs, *i.e.*, single cell output followed by patient output, we implemented ClassCox, ClassClass, ClassBlank, BlankClass, and BlankCox models based on equations (7)-(11) in the current version but intend to provide more options in the future.

To keep the analyses consistent, we used the same network architecture and hyperparameters throughout all of the experiments. Specifically, we used a three-layer DenseNet architecture bootstrap aggregated five times such that Eq. 1 would consist of a DenseNet instead of a single layer feedforward network and five such models were trained. The same set of hyper-parameters were used in all of the experiments in this study, except for the robustness to hyper-parameters experiment, where they were intentionally altered to test the influences on the output results. These are considered the default hyper-parameters in the *DEGAS* package but can be changed. They are: training steps 2000, single cell batch size 200, patient batch size 50, hidden layer nodes 50, drop-out retention rate 50%, single cell loss weight (*λ*_0_) 2, patient loss weight (*λ*_1_) 3, MMD loss weight (*λ*_2_) 3, and *L_2_*-regularization weight (*λ*_3_) 3.

### Feature selection and scaling

There are already multiple feature selection techniques available in a wide range of general statistical packages and scRNA-seq packages. For this reason, *DEGAS* does not focus mainly on feature selection, data cleaning, scRNA-seq clustering, but rather on transferring clinical traits from patient to cells for the purpose of prioritizing those cells. For these reasons, a wide range of feature selection techniques can be used before the *DEGAS* framework is applied.

Data from scRNA-seq experiments are generally very sparse. As a result, there are few genes with viable expression for any given cell. Due to this, it is necessary to perform feature selection to remove genes that are lowly expressed or have very low variance. When we select for high variance and expressed genes in the bulk expression data, more genes are filtered out. After the intersection of these two gene sets of expressed and high variance genes, we are left with less than 1000 genes. It is worth noting that such number of gene features is comparable to Seurat analysis, when usually hundreds to a couple of thousand highly variable genes are selected. The feature selection steps were tailored to each dataset because the data sparsity and variance vary greatly from one another, thus the tailored selection insured that enough genes with high enough variability were available to train on. The feature selection steps are described individually in each of the simulated, GBM, AD, and MM experiment sections.

For each experiment, the final feature scaling steps were consistent. The gene expression was converted to sample-wise z-scores because it allows the genes to be more comparable between samples and has been performed in multiple other studies [27, 48–50]. As the input to our deep learning models, we scaled these z-scores to a range of [0, 1]. This form of z-score scaling and [0, 1] scaling is commonly used in machine learning and deep learning to help model training [51–53]. We follow this same convention for our deep learning models.

### Disease association scores

The final *DEGAS* output is either the output of a sigmoid or a softmax activation. For these reasons, it can be useful to convert the [0, 1] label output to an association score which can be interpreted like a correlation coefficient. For these reasons, the output probability matrix from *DEGAS* can be converted to a [-1,1] value using the *toCorrCoeff* function in the *DEGAS* package. This function transforms the [0, 1] output value matrix (*P*) with *k* labels to [-1,1] using Eq. 12.

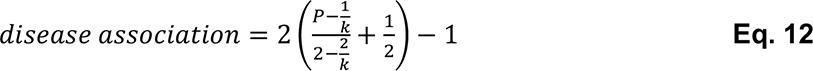

### Validating DEGAS using Simulated single cell data

First we generated 5,000 single cells in four cell types where the cell type 4 had two subtypes (cell type 4 disease and cell type 4 normal). Each of these five groups described above contains 1,000 cells. We split randomly these cells into 2 parts with 2,000 cells used for patient bulk tissue data generation and 3,000 cells to use directly as single cell data. The 2,000 single cells used to generate 600 patients across three different experiments (designated as simulation 1, 2, and 3) where in simulation 1 the cell type 1 is associated with disease, in simulation 2 only the cell type 4 disease is associated with disease, and in simulation 3 the entire cell type 4 is associated with disease. Each patient bulk tissue data was generated by randomly combining 400 single cells using the proportions in Table 3.

**Table 3.** Patient cellular makeup for simulation experiments. The abbreviations are: Simulation (sim), Normal (N), and Disease (D). The high-risk cell types are in bold.

|  | **Cell type 1** | **Cell type 2** | **Cell type 3** | **Cell type 4N** | **Cell type 4D** |
| --- | --- | --- | --- | --- | --- |
| Patients sim1D | **50.0%** | 16.6% | 16.6% | 16.6% | 00.0% |
| Patients sim1N | 25.0% | 25.0% | 25.0% | 25.0% | 00.0% |
| Patients sim2D | 25.0% | 25.0% | 25.0% | 00.0% | **25.0%** |
| Patients sim2N | 25.0% | 25.0% | 25.0% | 25.0% | 00.0% |
| Patients sim3D | 16.6% | 16.6% | 16.6% | **30.0%** | **20.0%** |
| Patients sim3N | 25.0% | 25.0% | 25.0% | 25.0% | 00.0% |

We then performed 10-fold cross validation by training the *DEGAS* ClassClass models using cell type and disease attribute. A total of 1000 gene features were used during training. We evaluated the model capacity for mapping patient labels on patients and cell type labels on single cells using PR-AUC and ROC-AUC. We then recapitulated the known cell type associations in each simulation by overlaying disease association onto the simulated cells. As a comparison, we also deconvoluted the patients using the 4 cell types using least squares. Deconvolution should be able to correctly identify the cells of interest in simulation 1 and simulation 3. In contrast, cell type prioritization using *Augur* [19] should be able to correctly identify the disease associated cell types in simulation 2. In the simulation 1 *Augur* experiment, cell type 1, cell type 2, cell type 3, and cell type 4 normal were randomly assigned to the disease or normal groups. In the simulation 2 *Augur* experiment, cell type 1, cell type 2, and cell type 3 were randomly assigned to disease or normal groups. The cell type 4 disease cells were all assigned to the disease group and the cell type 4 normal cells were all assigned to the normal group. In the simulation 3 *Augur* experiment, cell type 1, cell type 2, and cell type 3 were randomly assigned to disease and normal groups. The cell type 4 cells assigned to the disease group consisted of 60% cell type 4 normal and 40% cell type 4 disease and cell type 4 cells assigned to the normal group consisted of 100% cell type 4 normal. These cell type proportions match those in the simulation 3 patients used by *DEGAS*. The *Augur* output for each cell type is an ROC-AUC score that reflects how much a cell type changes transcriptionally between disease and normal samples. To make the comparison fair between our two methods, we use the output of our algorithm scaled from [0, 1] where 0.5 implies no association, 0 implies a negative association, and 1 implies a positive association. ROC-AUC is on the same scale. In this way we compare the strength of signal between *Augur* and our method to identify that cell type 4 has cell-intrinsic changes related to disease.

### Validating DEGAS using GBM data

The scRNA-seq data from the Patel *et al.* study [44] were downloaded from NCBI Gene Expression Omnibus (GSE57872). The single cell expression values were previously normalized to TPM containing 5,948 genes with *mean*(log_2_(TMP))>4.5 retained in the data table. The top 20% variance genes were retained for training. These values were converted to z-scores then standardized to a range of [0, 1] for each sample. The TCGA GBM microarray expression data was downloaded from *Firebrowse* (http://firebrowse.org/). Microarray data were used since it contains more patient samples for training with GBM subtype information than RNA-seq data. Likewise, the top 20% variance genes were retained for training and these expression values were converted to z-scores then standardized to a range of [0, 1] for each sample. The GBM subtype labels for the TCGA patients were downloaded from Verhaak *et al.* [54]. The intersection of genes between single cells and patients (199 genes) were used for the final model training. Since subtype labels were only available for the GBM patient samples, we trained a BlankClass *DEGAS* model (Eq. 10). This model minimizes the MMD loss between single cells and patients while minimizing the classification loss only in GBM patients. We split the dataset into 10 groups and performed 10-fold cross-validation by leaving out a single patient group during training. After cross-validation, we converted the [0, 1] *DEGAS* output to an association [-1,1] using the *DEGAS toCorrCoeff* function. These association scores were overlaid on the GBM single cells and now referred to as GBM subtype association scores because GBM subtype from patients is overlaid on single cells. We plotted these association scores stratified by GBM subtype for each tumor individually. We then compared the proportions of these cell types to the previously defined GBM types from the original publication were marked with red boxes. We also visualized the GBM subtypes association in single cells by calculating a low dimensional representation using *tSNE* and overlaying the *kNN* smoothed GBM subtype associations. To make the scatter plots of cells and patients more informative, *kNN* smoothing was used by averaging each point’s GBM subtype association value with its five nearest neighbors in *tSNE*. The model performance was shown with the PR-AUC and ROC-AUC for each of the GBM subtype labels in the TCGA patients from cross-validation.

In a second analysis on the GBM scRNA-seq and bulk expression data, using the same input features, we overlaid risk derived from the overall survival in the TCGA GBM cohort onto the individual cells from the Patel et al. study [44]. GBM has an extremely low 5-year survival rate resulting only three patients being censored. We introduced more censoring in the data by generating a uniformly distributed random vector of censoring times in the range 1 to 1063 days, where 1063 days is the 90th percentile of survival times. If the censor time was lower than the survival time, the patient was censored at that time instead of having an event at their true survival time. We then trained 10 BlankCox *DEGAS* models based on the patient survival input during 10-fold cross validation. The output from these *DEGAS* models were *kNN* smoothed based on the *tSNE* coordinates using the *DEGAS knnSmooth* function and converted to death associations using the *DEGAS toCorrCoeff* function. To highlight the differences in death association of cells, these associations were centered to 0 using the *DEGAS centerFunc* function. We evaluated the accuracy of the labels in patients using a rank-sum test based on the cox output in the GBM patients.

### Validating DEGAS and exploration using AD data

For AD datasets, we were primarily interested in identifying known relationships between cell types and AD diagnosis. For these reasons, we downloaded all of the adult Human scRNA-seq data from the AIBS. Only inhibitory neurons, excitatory neurons, oligodendrocytes, astrocytes, microglia, and oligodendrocyte progenitor cells (OPCs) were retained in the analysis due to the extremely low sample sizes for the remaining cell types. The inhibitory and excitatory neuron groups were merged into a single neuron group. These data were then log_2_ transformed, converted to sample-wise z-scores, and then standardized to [0, 1] by each sample. In the primary analysis, only the top 50 up-regulated DEGs for each cell type (calculated by *Seurat*) were retained in the single cell data (see **RESULTS**). In a distinct secondary analysis, features were selected with >25% non-zero samples and top 20% variance genes (see **Supplementary Materials**). The labels for the single cells consisted of the major cell types listed above. The AD brain data was downloaded from Mount Sinai/JJ Peters VA Medical Center Brain Bank (https://www.synapse.org/#!Synapse:syn3157743). Each of the RNA-seq samples were either from an AD patient’s brain sample or a normal control brain sample. The binary disease attribute of AD case or normal were used as the label for the model. Like in the previous experiment, the RNA-seq values were log_2_ transformed, converted to sample-wise z-scores, and standardized to [0, 1] for each sample. The top 50% variance genes were retained for training to keep the feature set larger. The intersection of the patient genes and single cell genes (Primary analysis: 169 genes, Secondary analysis: 456 genes) were using to train the final models. Using the cell type classification for each AIBS single cell and the AD/normal classification for each MSBB patient we were able to train a *DEGAS* ClassClass model (Eq. 8). The performance was evaluated using 10-fold cross-validation by leaving out each group during training once. As in the GBM experiments, we converted the *DEGAS* output to an association using the *DEGAS toCorrCoeff* function for each single cell so that each single cell now had an AD association. Correlation analysis was performed on AD association scores for different cells with each cell type by taking the median score and calculating the p-value by treating it as a correlation. In addition, single cells were plotted overlaid with *kNN* smoothed AD association. Furthermore, to evaluate *DEGAS* performance, PR-AUC and ROC-AUC were computed for the single cells during cross-validation for each cell type in the single cell data. Similarly, AD diagnosis PR-AUC and ROC-AUC were computed from the MSBB patient RNA-seq. For both the primary and secondary AIBS analysis, DEGs were identified for the high AD association astrocytes and microglia based on the median AD association then compared to their respective disease associated astrocyte (DAA) [55] (**Supplementary File 1**), human Alzheimer’s microglia (HAM) gene markers [56] (**Supplementary File 2**), or disease associated microglia (DAM) gene markers [57] (**Supplementary File 3**). A detailed description of these gene lists can be found in the **Supplementary Materials DAA, HAM, and DAM markers** section.

To further highlight the cellular associations to AD, we also performed experiments using a scRNA-seq dataset from Grubman *et al.* [32]. Since this dataset was sparser, genes were used with >25% non-zero samples then the top 50% variance genes were selected from these. For the MSBB data, the same initial feature selection was used (top 50% variance). The same normalization and standardization procedure as the AIBS scRNA-seq and MSBB were used again. The intersecting genes between Grubman *et al.* scRNA-seq constituted the final feature set (61 genes). 10-fold cross validation was performed using a ClassClass model and the AD associations were overlaid onto the Grubman *et al.* scRNA-seq in the same fashion as the previous experiment. In addition, a targeted analysis on only the microglia cells was performed. A single BlankClass model was trained using the same 61 features on the entire Grubman *et al.* microglia scRNA-seq and MSBB RNA-seq. For both analyses, the AD associations were overlaid onto the cells, AD associations were compared between cells from AD and normal patient samples, and DEGs were identified for the high AD association astrocytes and microglia based on the median AD association then compared to their respective DAA [55] (**Supplementary File 1**), HAM gene markers [56] (**Supplementary File 2**), or DAM gene markers [57] (**Supplementary File 3**). For the targeted analysis on only microglia, correlation tests were performed between AD associations and HAM gene markers [56] (**Supplementary File 2**). Also, DEGs were identified for the high AD association microglia based on the median AD association then compared to the HAM gene markers [56] (**Supplementary File 2**) and DAM gene markers [57] (**Supplementary File 3**).

Lastly, *DEGAS* analysis was performed on the Mathys *et al.* scRNA-seq dataset [15]. In this analysis, the same gene set as the AIBS Primary analysis, *i.e.*, all overlapping genes (157 genes) were used as input features. 1000 cells or all cells if total number was less than 1000 were sampled from each cell type since some cell types were over-represented. The same normalization and standardization procedure was used as the previous analyses. 10-fold cross validation was performed using these cells from Mathys et al. and the MSBB patient RNA-seq data using cell type and patient AD status as outcomes respectively. These outcomes represent a ClassClass DEGAS model. From the cross-validation results, the ROC-AUCs and PR-AUCs for each cell type label and the patient AD status were calculated. AD associations were calculated in the same fashion as all previous analyses. The Disease associations were then compared with AD status of the scRNA-seq donors and across the cell types. DEGs were identified for the high AD association astrocytes and microglia based on the median AD association then compared to their respective DAA [55] (**Supplementary File 1**), HAM gene markers [56] (**Supplementary File 2**), or DAM gene markers [57] (**Supplementary File 3**).

### Preprocessing of MM scRNA-seq

The scRNA-seq data were first combined into a dataset using *Seurat-CCA* [28]. This initial dataset integration allowed conserved subtypes of cells to be identified across datasets. All four patient dataset counts were loaded into a *Seurat* object. *Seurat* normalized, scaled, removed poor quality cells, and identified high variance genes. Using the union of high variance genes, multi-canonical correlation analysis was run across all four datasets, the subspaces were aligned across patients, the aligned single cells were plotted with *tSNE* [58], and clusters of cells were identified. The raw expression values for the high variance genes identified by *Seurat* were log_2_ transformed, converted to z-scores, and then scaled to [0, 1].

Furthermore, each IUSM scRNA-seq patient was individually clustered using *Seurat* to check the replicability of the clusters and were plotted with *UMAP* [59]. We used Rand, Fowlkes and Mallows’s index (FM), and Jaccard index (JI) to measure the cluster consistency between single patient clustering experiments and the merged all-patient clustering results. The four single patient clustering results, one for each IUSM scRNA-seq patient, were used as input into *BERMUDA* [25] to visualize and evaluate the original Seurat clustering.

### Preprocessing of MMRF patient data

MMRF patients with bulk tissue RNA-seq and clinical data were used in MM analysis. We used PFS as the disease attribute of interest. TPM values for the MMRF patient gene expression data and the PFS data were used as the input for *DEGAS*, these values were log_2_ transformed, converted to z-scores, and scaled to [0, 1]. The union of the features (502 genes) identified by Seurat in the single cell data and the features selected in the MMRF patient data were used as the final feature set. The features retained in the MMRF data were identified by fitting an elastic-net Cox model [60] to the TPM values based on the PFS.

### Evaluate DEGAS performance on MM datasets

PR-AUC and AUC were calculated for each of the output labels for the single cells and for patient labels if a classification output was used for the patient data. Cox proportional hazard output was used on patients, a log-rank test was calculated for each patient so that the hazard ratio and p-value could be evaluated based on patient stratification by median proportional hazard. Additionally, the same models were used to predict risk in the GSE2658 dataset which had information on OS. The output for each GSE2658 sample averaged across all 10 *DEGAS* models and stratified by median risk to show the robustness of the cox output across datasets.

### Identification of CD138+ cell types associated with MM prognosis

The single cells from MM patients can be assigned proportional hazards based on the MMRF Cox output of the model. Each single cell in the validation set was assigned progression association by feeding those samples through the Cox output layer. In this way, we can infer the association with progression risk of specific cell types as well as the cell type enrichment contained in each MMRF sample. Since the Cox output is a proportional hazard, we centered the outputs to zero for each step of cross validation to produce a PFS association using the *DEGAS centerFunc*. We plotted these relationships and conducted Student’s t-tests on the subtype vs. PFS association in IUSM single cells, PFS association vs. MM malignancy from Ledergor *et al.*, and subtype 2 enrichment vs. MM malignancy from Ledergor *et al* [42].

### Analysis of differential gene expression in prognostic cell types

T-tests were calculated cell subtype 1 vs all cell subtypes and cell subtype 2 vs. all cell subtypes using the batch corrected gene expression values from *Seurat*. These values were stored in (**Supplementary File 4** and **Supplementary File 5**) respectively. For the marker set of *PHF19*, *HELLS*, *EZH2*, *TYMS*, *ZWINT*, and *MKI67* we performed t-tests for each patient individually.

### Evaluation of DEGAS robustness to hyper-parameters in GBM

Using the GBM dataset, we evaluated the robustness of DEGAS model outputs to hyper-parameters by repeating 10-fold cross-validation 100 times with randomly generated hyper-parameters following a uniform distribution. The range of hyper-parameters used in training consisted of training steps 1,000-3,000, single cell batch size 100-300, patient batch size 20-100, hidden features 10-100, drop-out retention rate 0.1-0.9, Cell loss weight (*λ*_0_) held at constant 2, Patient loss weight (*λ*_1_) 0.2-5, MMD loss weight (*λ*_2_) 0.2-5, *L_2_*-regularization weight (*λ*_3_) 0.2-5.

Using these outputs, we performed two tests. One was to evaluate the loss in performance based on changing the hyper-parameters where performance was measured with ROC-AUC among the TCGA GBM patients labeled by patient GBM subtype (Mesenchymal, Classical, Proneural, Neural). In this test, we calculated the spearman correlation and plotted the scatter plot between the AUC of each of the four GBM subtype labels and the hyper-parameters used.

Next, we evaluated whether or not the correct GBM subtype labels (Mesenchymal, Classical, Proneural, Neural) could be recapitulated in the GBM scRNA-seq tumors that had known GBM subtypes (MGH26: Proneural, MGH28: Mesenchymal, MGH29: Mesenchymal, MGH30: Classical). To do this for each tumor (MGH26, MGH28, MGH29, MGH30), the rank of the correct label was calculated by calculating the mean of each GBM subtype association across all of the cells in that tumor. This resulted in each of the 100 random hyper-parameters having a rank for each GBM subtype for each of the GBM scRNA-seq tumors (4 highest ranked, 1 lowest ranked). Ideally all GBM scRNA-seq tumors would have a rank of 4 indicating the correct GBM subtype was ranked the highest regardless of hyper-parameters. Similarly, we also calculated the spearman correlation and plotted the scatter plot between correct label rank and the hyper-parameters used.

### Evaluation of domain adaptation for DEGAS disease association transfer

We evaluate the necessity for domain adaptation to transfer disease associations to single cells using 30 total experiments. These experiments evaluated disease associations in cells by training with MMD loss vs. those without MMD loss for a variety of biases added between the cells and patients. It is important to highlight the fact that without bias between different datasets, in this case cells and patients, there is no need for domain adaptation. Practically in real transcriptomic data, there will always be bias between datasets. For these reasons we added bias for these 30 experiments. These experiments were conducted for every combination of MMD loss (with and without MMD), simulation (three simulations), and cellular subtype (five total subtypes since cell type 4 has two subtypes) totaling 30 combinations. The experiments were conducted as follows. In each experiment, the counts of 300 cells from a given subtype were aggregated together and multiplied by 1000 constituting a large systematic bias associated with a single subtype. This bias vector was added to all of the patients in the given simulation, both disease and normal. A single three-layer DenseNet *DEGAS* model with five-fold bootstrap aggregation was trained on all the cells and all the patients then the disease associations were predicted in the cells. We evaluated error by subtracting the expected disease association from the predicted disease associations, *e.g.*, cell type 1 in simulation 1 should be 1. We then compared the error rates between the *DEGAS* models with and without MMD using a t-test.

### Evaluation of regularization in DEGAS performance

Regularization is an important method in machine learning to prevent model overfitting. Here we utilized three such techniques to prevent overfitting, namely, *L_2_*-regularization, dropout, and bootstrap aggregation. Since all of these techniques may work better or worse in different scenarios, we perform a simple experiment where all of these regularization techniques are removed and compared with the regularized results. We performed experiments using each of the simulated datasets. To evaluate the robustness of our models we performed 10-fold cross validation in each simulation. The simulated cells were split into 10 groups and the simulated patients were split into 10 groups. For each fold of cross validation, our default *DEGAS* three-layer DenseNet model with *L_2_*-regularization, dropout, and bootstrap aggregated 5 times was trained then a three-layer DenseNet *DEGAS* model was trained on the same data without *L_2_*-regularization, dropout, and bootstrap aggregation. Both models were then used to predict the patient disease attributes in the holdout group of patients, the cell types in the hold out group of cells, and the patient disease attributes in the cells. We compare the performances using ROC-AUC and PR-AUC for patient disease status in patients and cell type in cells. Furthermore, we evaluate the label transfer of patient labels to cells by calculating the error based on the expected cell type association for each cell. We compare between the regularized and unregularized error in cells with a t-test.

## Results

### DEGAS clinical impression framework

In this study, we applied *DEGAS* to integrate and analyze scRNA-seq, bulk gene expression, and clinical data (Fig. 1) from simulated data as well as three different diseases: GBM, AD, and MM. The simulated, GBM, and AD datasets primarily served as validation to demonstrate the feasibility and universality of the *DEGAS* transfer learning approach since the ground truth of the simulated data was known, the correct GBM subtypes were known, and neuron loss with microglia gain in AD brains were also known. We then further expand our study to MM data, which serves as the discovery dataset, since the myeloma cell subtypes and high-risk factors related to MM are not as well understood at the single-cell level. In the MM study, we applied *DEGAS* on patient data from the Multiple Myeloma Research Foundation CoMMpass study (MMRF) and scRNA-seq data that we generated from myeloma patients. Our aim was to identify the cell subtypes using the impressions of progression risk on the single cells. We then applied the results to two separate MM validation datasets, one of which contained plasma cells from normal bone marrow (NHIP), two MM precursor conditions - monoclonal gammopathy of undetermined significance (MGUS) and smoldering multiple myeloma (SMM), and MM. We tested if *DEGAS* assignment of progression risk to cell subtypes were higher for more malignant conditions. An additional external validation dataset of patient level expression data with OS was used to evaluate whether the patient stratification learned by *DEGAS* was robust enough to be generalized to an external survival dataset.

### DEGAS correctly identifies high-risk cell types and subtypes in simulated data

To evaluate *DEGAS* in a controlled context, 5,000 single cells were generated with *Splatter* [61] (Fig. 2A) where 2,000 of the cells were held-out to generate simulated patients. Using this group of held-out cells, 600 simulated patients were generated by aggregating sets of 400 simulated cells (Fig. 2B-D). We conducted three simulation experiments, denoted Simulation 1, Simulation 2, and Simulation 3, where the single cells were aggregated in known proportions for each patient so that we could generate a “disease” patient group with different cellular composition than the “normal” patient group (**see Methods**). To highlight the utility of *DEGAS*, the experiments were: Simulation 1: cell type 1 is enriched in disease patients (Fig. 2B); Simulation 2: one subtype of cell type 4, *i.e.*, cell type 4 disease, is enriched in disease patients (Fig. 2C); and Simulation 3: both subtypes of cell type 4 are enriched in disease patients (Fig. 2D).

**Fig. 2.**
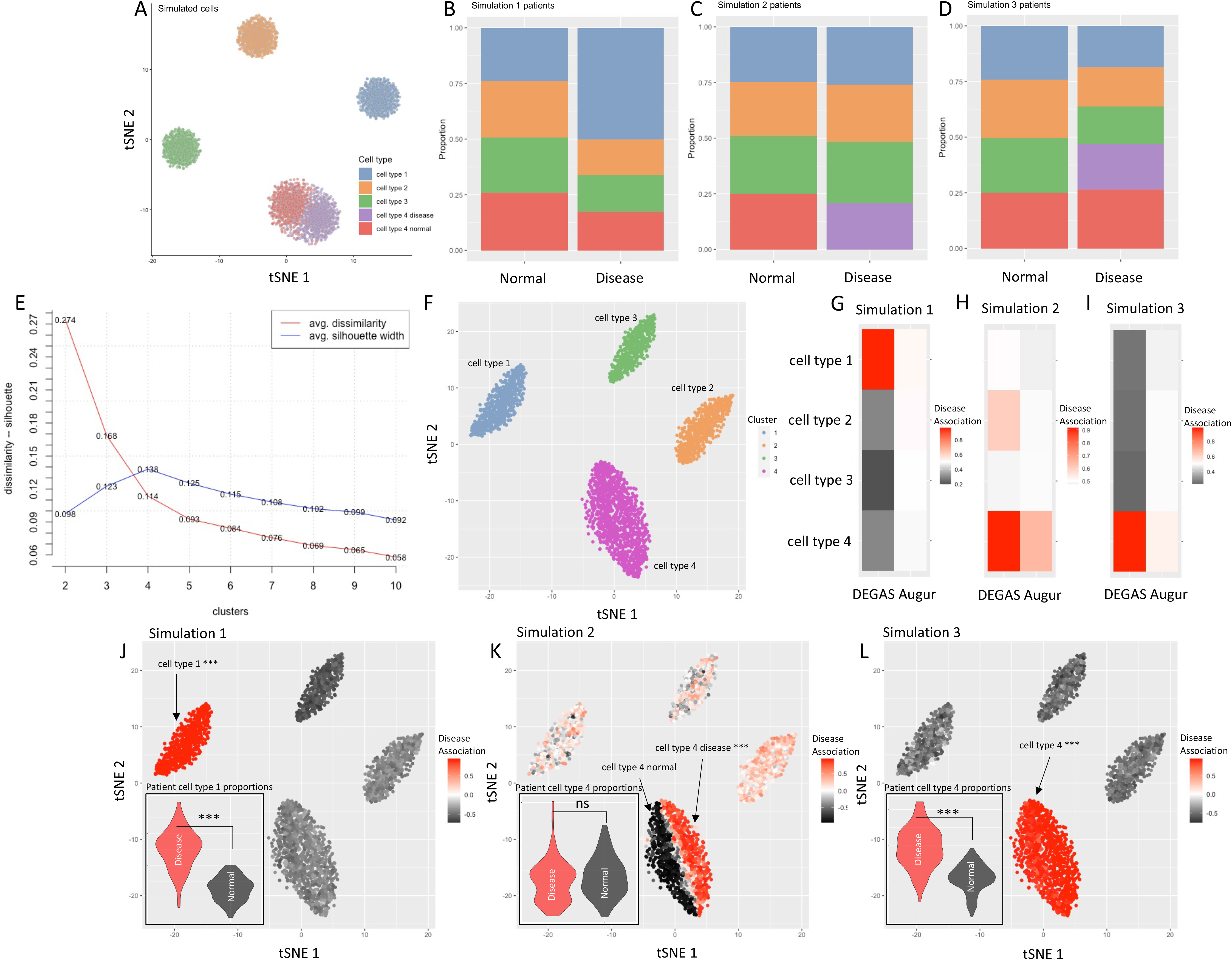
Simulation study and baseline comparisons of *DEGAS* framework. **A)** 5,000 simulated cells from *Splatter* with 4 cell types where one of the cell types has two subtypes. Cell type 4 is composed of two subtypes that are specific to either disease or normal patients. 2,000 of these cells were used to generate the 600 simulated patients in **B-D** and 3,000 were used as the cell input to our *DEGAS* models. **E)** Optimal cluster number (4 clusters) based on average silhouette width for the 3,000 cells not used to generate patients. **F)** The same 3,000 cells used as the cellular input colored by their cluster. **G)** *DEGAS* comparison to *Augur* in simulation 1. **H)** *DEGAS* comparison with *Augur* in simulation 2. **I)** *DEGAS* comparison with *Augur* in simulation 3. **J-L)** *DEGAS*-calculated disease association from each simulation overlaid onto 3,000 cells. The violin plot in the bottom left corner is deconvolution cell type proportion for cell type 1 in simulation 1 patients (**J**), cell type 4 proportion in simulation 2 patients (**K**), and cell type 4 proportion in simulation 3 patients (**L**).

Please note that the optimal number of clusters for the simulated single cells would be determined to be four based on a standard scRNA-seq workflow (*i.e., tSNE* followed by *K-Medoids* where optimal cluster number is selected based on average silhouette width) (Fig. 2E). This would cluster the cells into the four cell types while ignoring the two subtypes in the cell type 4 (Fig. 2F). As a result, deconvolution algorithms will not be able to detect the subtype level risk associations. Fortunately, cell type prioritization algorithms like *Augur* can detect these changes within cell types due to disease. However, for situations that do not have a new cell type or missing cell type in the disease (simulation 1), *Augur* cannot detect the association between cell type 1 and disease since there is no disease associated cell type change (Fig. 2G). *Augur* can detect the disease-associated cell type 4 in simulation 2 (Fig. 2H). In simulation 3 where there is a mix of disease and normal subtypes for the cell type 4 in the disease group, *Augur* again has difficulty in identifying the cell type 4 disease association (Fig. 2I). In contrast to *Augur*, deconvolution can easily identify the correct cell type for Simulation 1 (Fig. 2J) and Simulation 3 (Fig. 2L) but not for simulation 2 (Fig. 2K). In comparison, *DEGAS* not only identified the correct cell type and subtypes in each experiment, it also correctly detected all of the simulated disease associations (Fig. 2G-L). Additionally, *DEGAS* had high precision-recall area under the curve (PR-AUC) predicting disease status of simulated patients (0.96-0.98) (**Table S1**) and almost perfectly predicted the cell type of simulated cells (∼1.0) during cross-validation (**Table S2**). Since *DEGAS* directly assigns disease risk to cells, many of the problems with cell type level analyses can be avoided and the correct groups of cells can be identified by overlaying impressions of disease risk.

### DEGAS correctly mapped single cells to corresponding GBM subtypes

We first demonstrate *DEGAS* in a straightforward case to show the performance of our framework using real data from GBM. We use single-cell data from Patel *et al.* [44], in which researchers assigned four major GBM tumor subtypes (Proneural, Mesenchymal, Classical, and Neural) to the scRNA-seq data obtained from five GBM tumors. Of the five tumor samples, four had been labeled in the original publication with a single subtype based on the major proportion of cells assigned to each GBM subtype. For GBM bulk tumor tissue expression data, we obtained microarray data for 111 GBM patients from The Cancer Genome Atlas (TCGA), for which the same labels of GBM subtypes were also provided. The OS was also available in a subset of 109 patients. As the simplest form of validation, we used these two datasets as input for the *DEGAS* model to test if it could re-identify the same GBM subtypes for both single cells and the TCGA GBM cohort simultaneously. Then we overlaid OS-derived death associations onto the cells to visualize their association with OS. The resulting *DEGAS* models also proved to be accurate with high PR-AUCs (0.79-0.97) when predicting each of the GBM subtypes in the TCGA patients during 10-fold cross validation (**Table S3**). The OS BlankCox *DEGAS* models were able to stratify the patients into high and low risk groups based on median patient risk (log-rank p-value < 0.05). *DEGAS* correctly re-identified the same labels for all four tumors by overlaying GBM subtypes associations on each single cell, as indicated by the groups of cell subtypes with the highest association score determined by the median value (indicated with a red box) (Fig. 3A-D). For the fifth tumor sample, MGH31, it was labeled as a combination of multiple GBM subtypes in the original study, so we did not use it in our evaluation although *DEGAS* identified mesenchymal as its most associated GBM subtype (Fig. 3E). Additionally, these relationships can be visualized by plotting the single cells and overlaying the GBM subtype association or OS-derived death association. It is clear that MGH28 and MGH29 have a high association with the mesenchymal GBM subtype (**Fig. S1A**) and contain populations of cells with high death associations (Fig. 3F).

**Fig. 3.**
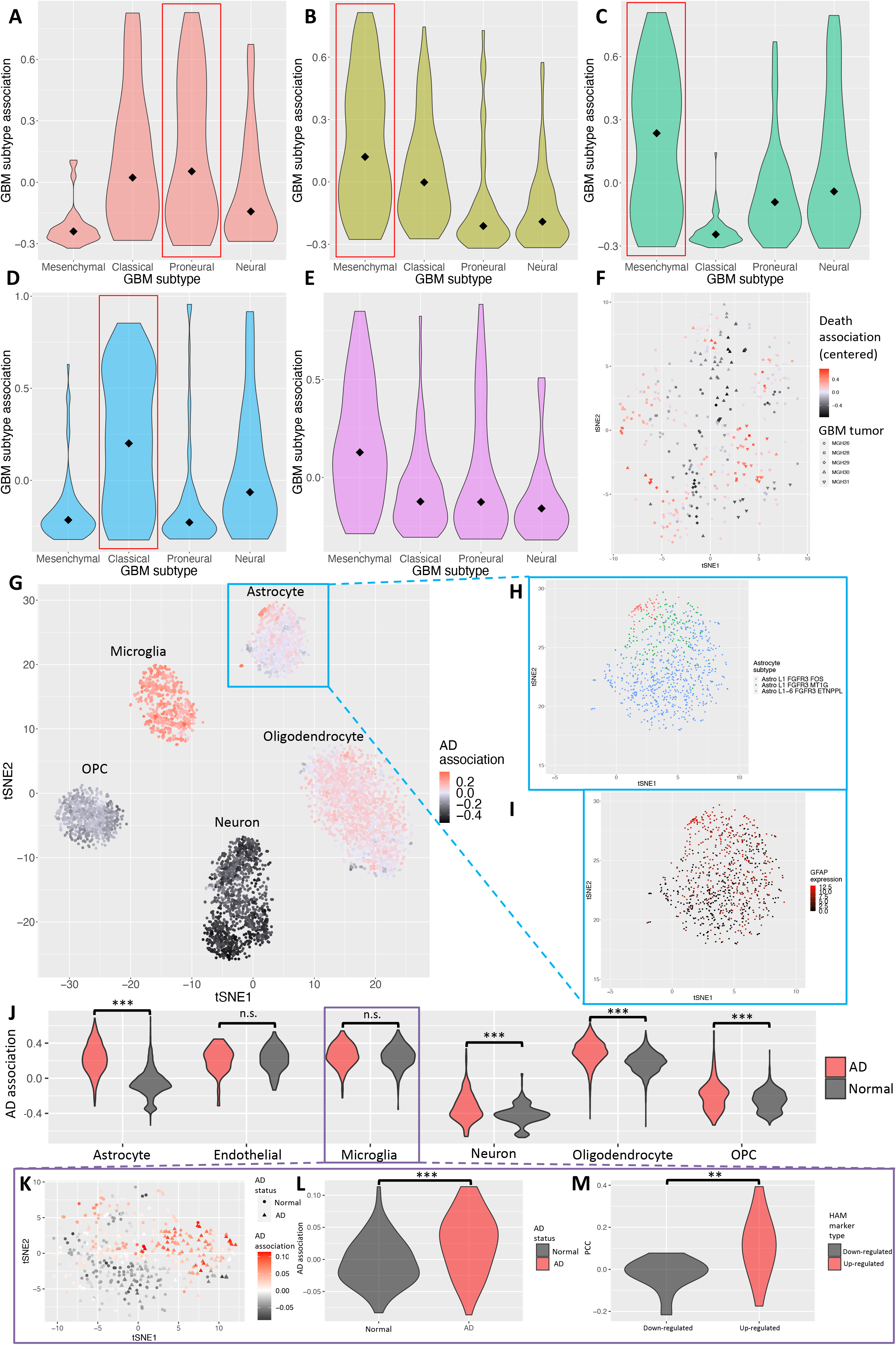
*DEGAS* validation in GBM and AD. *DEGAS* output of the distribution of GBM subtypes in single cells from five GBM tumors. Four of the five tumors had known GBM subtype information from Patel *et al.* (MGH26: Proneural, MGH28: Mesenchymal, MGH29: Mesenchymal, and MGH30: Classical, indicated by red boxes) which were recapitulated by *DEGAS*. The subtype information for the tumors, MGH26, MGH28, MGH29, and MGH30 were derived from Patel *et al.* where MGH31 did not have a clearly defined GBM subtype. The association of cells assigned to each subtype were plotted for each tumor; **A)** MGH26, **B)** MGH28, **C)** MGH29, **D)** MGH30 and **E)** MGH31. Median values are marked by a diamond in each of the violin plots. **F)** The death association centered around 0 is overlaid on all of the single cells from the five tumors (indicated by color). **G)** *DEGAS* output of AD association for each single cell. The AD association score is indicated by the color and is overlaid onto AIBS single cells. This plot shows the negative AD association in neuron cells and positive AD association in Microglia. **H-I)** There also appeared to be a subpopulation of astrocytes with positive AD association. The astrocytes were plotted separately and colored by AIBS Astrocyte subtypes **(H)** and *GFAP* expression, a disease-associated astrocyte marker **(I)**. **J)** Comparison of *DEGAS*-derived AD associations for single cells from AD and Normal control samples from Grubman *et al.* **K-M)** Targeted analysis of microglia from Grubman *et al.* including the AD associations overlaid onto microglia **(K)**, AD association comparing AD status of patient sample from which the cells were sampled **(L)**, and PCC between AD association with HAM marker genes comparing up-and down-regulated HAM marker genes **(M)**. Significance values: n.s. (not significant), • (0.1), * (0.05), ** (0.01), *** (0.001).

### DEGAS identifies increased microglia, reduced neuron populations, DAAs and DAMs

Aside from GBM, AD also has well documented characteristics that can be used as a test bed for *DEGAS*. Specifically, there is a well-documented reduction in neurons [36–38], increase in microglia [33–35, 39], and more recently, AD subtypes of astrocytes [55] and microglia [56, 57]. AD brain scRNA-seq data was obtained from the AIBS and bulk AD RNA-seq were retrieved from MSBB [46]. During 10-fold cross-validation, *DEGAS* models for both primary and secondary AIBS analyses achieved high AD diagnosis status PR-AUC (0.82 and 0.76) in MSBB patients **(Table S4**) and high cell type prediction PR-AUCs (>0.99) for AIBS single cells (**Table S5**).

From the AIBS primary analysis *DEGAS* results, we confirmed that at the single cell level, the AD associations were negative in neurons as previously described [62], which is shown by the dark shade of neurons compared to other cell types (Fig. 3G, Table 4). In contrast, we observed positive AD associations in microglia cells (Fig. 3G, Table 4). A strength of the *DEGAS* framework is that it can detect intra-cell type differences in disease risk. Within the astrocyte cell type, we identified an astrocyte subtype that had a positive association with AD (Fig. 3G) that corresponded to the *Astro L1 FGFR3 FOS* subtype from the AIBS brain cell atlas (*i.e., FOS* is a DAA marker) [63] (Fig. 3H), had up-regulated DAA marker *GFAP* (Fig. 3I) [55], and was enriched for DAA markers (OR = 30.93, Fisher’s exact p-value < 2.2•10^-16^**, Table S6**). Furthermore, the high AD association microglia were enriched for DAM markers (OR = 17.07, Fisher’s exact p-value = 2.11•10^-10^**, Table S7**). In the secondary AIBS analysis using high variance genes, we again identified the strong negative AD association for neurons (**Fig. S2A**), positive AD association in microglia (**Fig. S2A**), high AD association astrocytes enriched for DAA markers (OR = 5.65, Fisher’s exact p-value < 1.66•10^-8^**, Fig. S2B,C, Table S8**), and high AD association microglia enriched for DAM markers (OR = 14.34, Fisher’s exact p-value < 4.01•10^-11^**, Table S9**). When we performed *DEGAS* analysis on a separate dataset from Grubman et al. [32] with single cells from both AD and normal brains, we found that the major cell types from AD brains were significantly more associated with AD than their counterparts in normal brains as judged by median value (Fig. 3J).

**Table 4.**
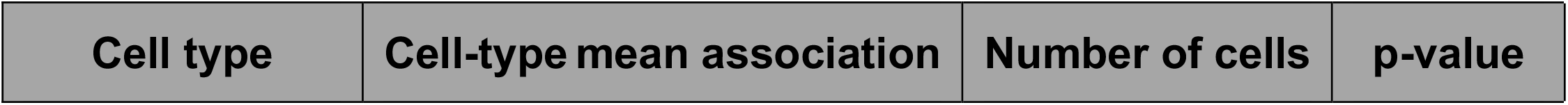
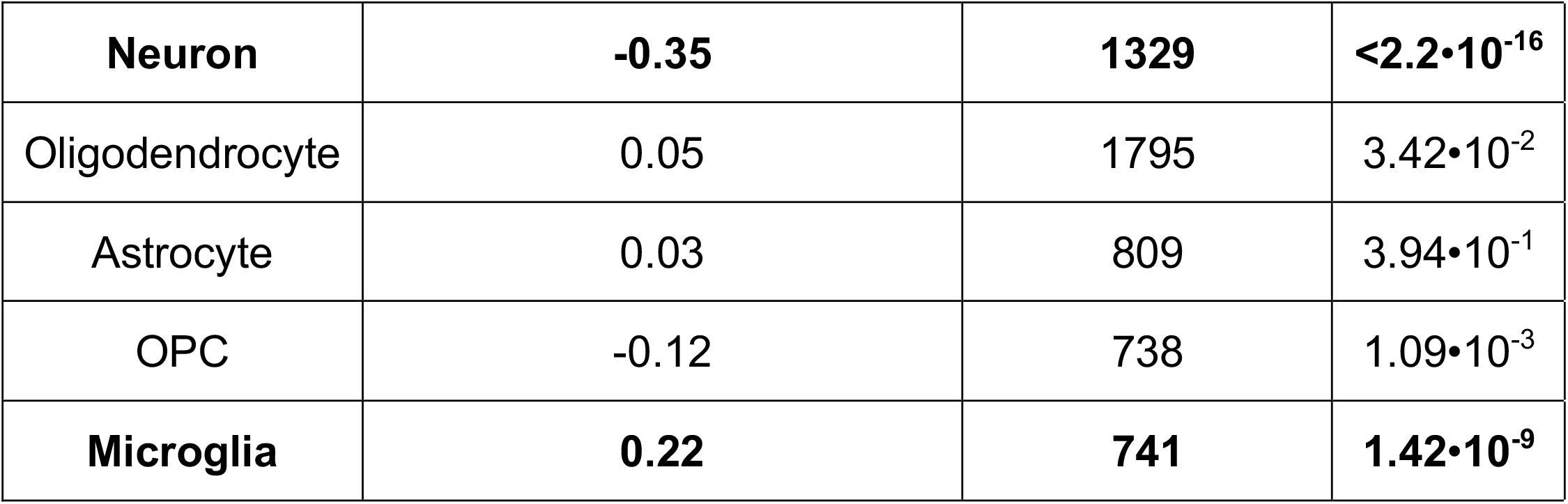
Comparison of AD association scores in single cells between cell types as visualized in Fig. 3G. The *DEGAS* models were trained using neuron, oligodendrocyte, astrocyte, OPC, and microglia cell types. The single cells were split into groups based on their cell type and the mean AD associations of each cell type was evaluated as a correlation. The neuron and microglia groups are bolded to highlight their much higher mean AD association. P-values are calculated by treating the association score as a pearson correlation coefficient.

In the Grubman *et al.* scRNA-seq data, the astrocytes in AD brains were highly positively associated with AD (AD association = 0.22, pearson correlation p-value = 7.89•10^-7^) whereas the astrocytes in normal brains were negatively associated with AD (AD association = −0.06, pearson correlation p-value = 1.06•10^-2^, Fig. 3J). Astrocytes from AD brains also expressed *GFAP* at greater levels than astrocytes from normal brains (t-test p-value < 2.20•10^-16^) and high AD association astrocytes were significantly enriched for DAA markers (OR = 21.90, Fishers exact p-value = 2.21•10^-12^, **Table S10**). Furthermore, the high AD association microglia were moderately enriched for DAM markers (OR = 4.15, Fishers exact p-value = 4.11•10^-2^, **Table S11**). This provides evidence for DAA and DAM cells in the Grubman *et al.* dataset.

DAM and HAM marker enriched high AD association cells were independently identified in the targeted analysis of the Grubman *et al.* microglia cells (Fig. 3K). AD associations were higher in cells derived from AD patient samples than Normal patient samples (Fig. 3L, t-test p-value = 6.66•10^-12^), HAM up-regulated markers were more likely to be significantly positively correlated to AD association than HAM down-regulated markers (Fig. 3M, t-test p-value = 2.63•10^-3^). The HAM marker APOE [56, 57] was positively correlated with AD association (**Table S12**, PCC=0.18, p-value = 1.15•10^-4^). High AD association microglia were significantly enriched for HAM markers (OR = 21.47, Fishers exact p-value = 6.33•10^-4^, **Table S13**) and DAM markers (OR = 11.52, Fishers exact p-value = 1.50•10^-11^, **Table S14**). It is important to note that there was no overlap between the input feature set used to train the DEGAS model and HAM marker genes that were identified, which shows DEGAS is a useful tool to identify disease associated cells within a single cell type even without prior knowledge of marker genes.

After applying *DEGAS* to the Mathys *et al.* scRNA-seq dataset, the DEGAS models achieved high AUCs for patient AD status (0.77), patient AD status PR-AUC (0.81), cell types (>0.98), as well as cell type PR-AUCs (0.82-0.99) during cross validation (**Table S15-16**). The positive AD association of microglia and negative AD association of neurons were recapitulated (**Fig. S3A, Table S17**). Within the astrocyte cluster, there existed a subset of astrocytes with higher AD association (**Fig. S3B**). High AD association astrocytes were significantly enriched for DAA markers (OR = 14.75, Fishers exact p-value = 3.16•10^-15^, **Table S18**). A closer comparison of the scRNA-seq revealed that the top 10% AD association astrocytes, had 2.5 times higher GFAP expression than the other astrocytes (t-test p-value = 6.36•10^-8^, **Fig. S3C**). In fact, like the Grubman *et al.* analysis, the AD association scores were higher in cells coming from AD patients than normal patients for every cell type in the Mathys *et al.* analysis (**Table S17**). Notably, we see increased AD association in AD derived astrocytes and microglia likely representing DAAs and HAMs respectively (**Table S17**). Furthermore, high AD association microglia were highly enriched for DAM markers (OR = 19.35, Fishers exact p-value < 2.2•10^-16^, **Table S19**) and high AD association in astrocytes correlated well with neuritic plaque count, a marker for disease severity in AD patients (PCC = 0.22, p-value = 6.36•10^-12^, **Table S17**). Again, the Mathys *et al.* analysis provides another example to demonstrate that DEGAS recapitulates the findings from the AIBS and Grubman *et al.* analyses and shows that DEGAS models can capture cell type level as well as intra-cell type differences in disease association.

### Identification of plasma cell subtypes in CD138+ scRNA-seq of MM

In the MM study, unlike the previous two datasets, there were no predefined cell type labels, but *DEGAS* was still capable of analyzing such data and give clinical perspective to the clusters of cells in the MM scRNA-seq data. In order to cluster cells into groups, we first used Seurat [28], a commonly used scRNA-seq data analysis tool, to merge and cluster all the CD138+ bone marrow cells from four patients (two SMM and two MM) whose samples were collected at the IUSM. Using *Seurat*, five major clusters of cells were identified (Fig. 4A). Cluster 1 consisted of the majority of the cells in each sample and was most likely the main clone in each of the patients. Cluster 2 was present in many of the patients and is described in detail after the *DEGAS* analysis. Cluster 3 and 5 were only present in patient 2 representing possible subclones in patient 2. Cluster 4 was shared between multiple patients. These five clusters were used as the subtype labels in the *DEGAS* framework. We verified these cell clusters by clustering cells from each patient individually with *Seurat* and another scRNA-seq normalization tool *BERMUDA* (Batch Effect ReMoval Using Deep Autoencoders) [25] for all four patients. We found that the individual clustering results closely mirrored the *Seurat-CCA* clusters (**Fig. S4A-D**, **Table S20**) and that the subtype 2 was consistent across all MM patients using *BERMUDA* (**Fig. S4E**). For bulk tissue data from MMRF, the clinical outcomes of PFS for 647 patients were used as the patient-level input to *DEGAS* and overlaid onto the CD138+ single cells from the four IUSM patients (Fig. 4B).

**Fig. 4.**
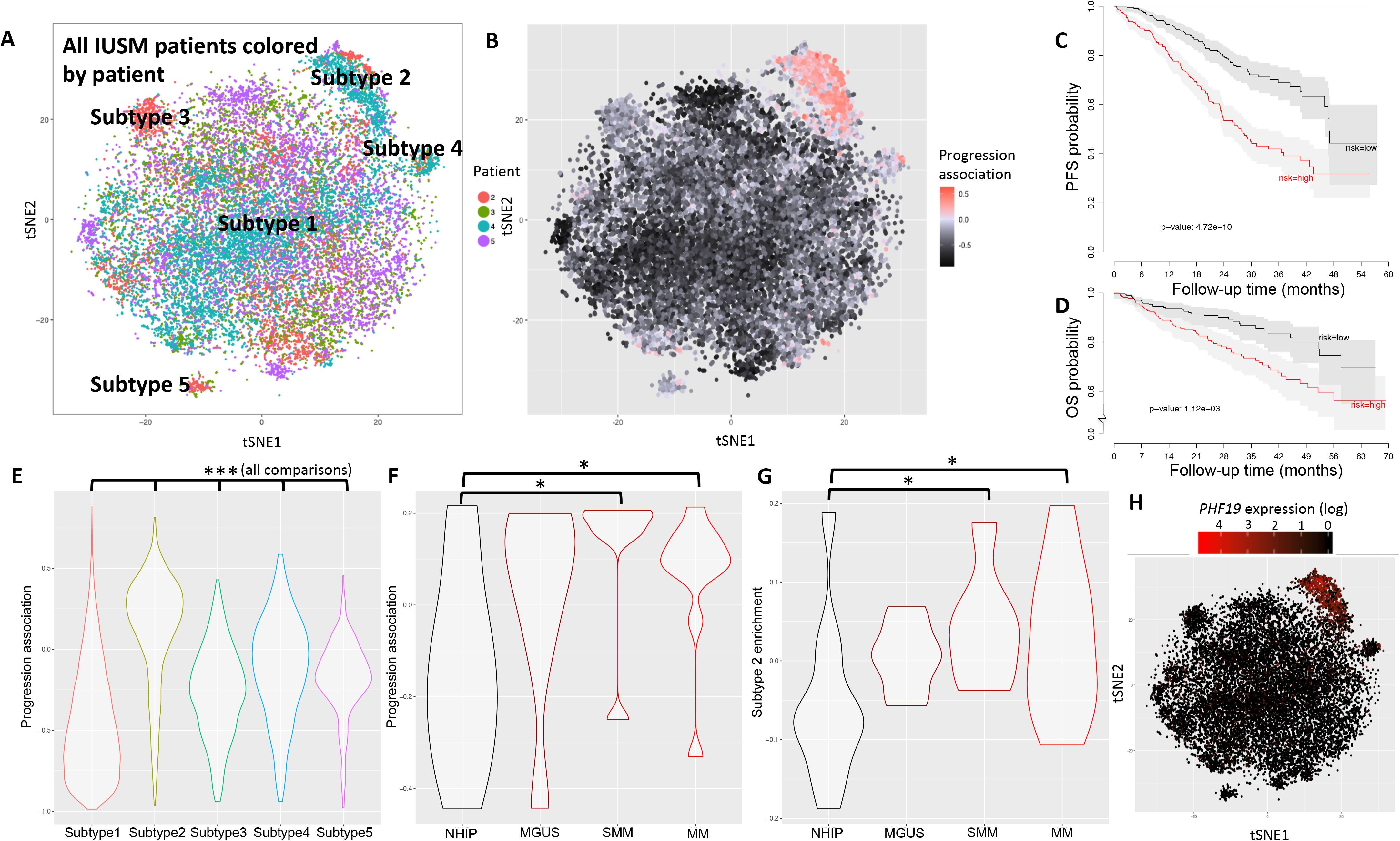
Association between subtypes and progression risk in MM. IUSM CD138+ scRNA-seq subtype clusters generated from *Seurat* colored by **A)** cluster, *i.e.*, subtype and **B)** progression association. **C)** Kaplan-Meier curves of PFS from cross-validation for the MMRF patients stratified by median proportional hazard. **D)** Kaplan-Meier curves of OS from Zhan *et al.* external dataset stratified by median proportional hazard. **E)** Progression association for IUSM CD138+ subtypes **F)** Progression association for NHIP, MGUS, SMM, and MM in the external dataset Ledergor *et al.* **G)** Subtype 2 enrichment for NHIP, MGUS, SMM, and MM in the external dataset Ledergor *et al.* NHIP: normal hip bone marrow, MGUS: monoclonal gammopathy of undetermined significance, SMM: smoldering multiple myeloma, MM: multiple myeloma. Significance values: • (0.1), * (0.05), ** (0.01), *** (0.001). All plots were generated using the default parameters for the *DEGAS* package described in the section of **Methods**: *Transfer learning using DEGAS*.

### DEGAS patient stratification and cell type classification on MM

A *DEGAS* model was trained on IUSM patient scRNA-seq data with subtype labels defined above and MMRF patients with bulk tissue data and PFS information. The performance metrics were calculated via 10-fold cross-validation. It is worth noting that for PR-AUC, random no skill classifiers will achieve a performance equal to the percentage of the class of interest and in the case of uncommon classes like subtype 4, the random classifier performance will be close to zero (0.02). When predicting cellular subtype label in single cells, *DEGAS* was able to achieve a PR-AUC between 0.44-0.98 for all of the five CD138+ cellular subtypes identified in the above scRNA-seq data while the PR-AUC for subtype 2 reached 0.91 (**Table S21**). The receiver operating curve AUCs (ROC-AUCs) were between 0.90-0.98 for these five subtypes (**Table S21**). Due to class imbalance some of the subtypes did not perform as well as others based on PR-AUC but all of the PR-AUCs were substantially greater than a purely random model. Aside from correctly classifying the single cells, *DEGAS* was able to stratify the MMRF patients into high and low risk groups based on median progression risk (log-rank p-value = 4.72•10^-10^, Fig. 4C). We then applied the trained model on an external patient transcriptomic dataset from Zhan *et al.* [41] for validation. We demonstrated that the Cox proportional hazards portion for patient OS time of the *DEGAS* model was robust across datasets, and the impressions extracted from the *DEGAS* framework were capable of stratifying patients into low-and high-risk groups in the validation dataset (log-rank p-value = 1.12•10^-3^, Fig. 4D).

### DEGAS identifies CD138+ cellular subtypes with high progression association

The MM scRNA-seq data provided an example of an exploratory analysis with *DEGAS* which can be used to generate hypotheses for future studies. The *DEGAS* model for the MM study transfers clinical impressions to single cells (*i.e.*, single cells were directly assigned a progression association score), as well as transfers cellular/molecular impressions to patients (*i.e.*, patients are assigned subtype enrichment score). We found that the subtype 2 cells were the most associated with prognosis (Fig. 4B) based on the *DEGAS* results. Specifically, the subtype 2 cells were associated with a shorter time to progression (Fig. 4E, t-test p-value < 2.2•10^-16^). On an external validation scRNA-seq dataset from Ledergor *et al.* [42], the progression association increased from NHIP (no disease) to SMM (Fig. 4F, t-test p-value = 1.50•10^-2^) and MM (Fig. 4F, t-test p-value = 1.70•10^-2^), which is consistent with the order of precursor conditions for MM (NHIP → MGUS → SMM → MM). In addition, the enrichment of the subtype 2 cells increased from NHIP to near-MM stage SMM (Fig. 4G, t-test p-value = 3.10•10^-2^) and MM (Fig. 4G, t-test p-value = 3.40•10^-2^).

### MM prognostic subtypes have distinct gene signatures

Differential gene expression analysis was performed between subtype 2 and all other subtypes (**Supplementary File 5**), and we found that subtype 2 had significantly up-regulated *PHF19* expression in all four of the patients (Fig. 4H). *PHF19* is a known marker for malignant disease in MM [64]. Besides *PHF19*, its associated markers such as *HELLS*, *EZH2*, *TYMS*, *ZWINT*, and *MKI67* were also significantly up-regulated in subtype 2. These results suggested the possible existence of a more malignant CD138+/*PHF19^high^* subpopulation of plasma cells represented by the subtype 2 cluster. It is important to notice that the gene feature set that was used as input into *DEGAS* only contained the *HELLS* gene, which further highlights the ability of *DEGAS* to predict high-risk cellular subtypes that can be further studied.

### DEGAS is robust to hyper-parameter choice

To assess the robustness of *DEGAS*, we also analyzed how the hyper-parameter choices influence its results using a set of 100 randomly generated hyper-parameters with 10-fold cross-validation on each set of those 100 sets of hyper-parameters on the GBM datasets. The hyper-parameters that we evaluated include: the number of training steps, batch size for single cells, batch size for patients, number of hidden layer nodes, drop-out retention rate (the percentage of nodes randomly retained at the hidden layer), patient loss weight, MMD loss weight, and *L_2_*-regularization weight. The detailed information about the range of hyper-parameters that were randomly sampled can be found in subsection titled *Evaluation of DEGAS robustness to hyper-parameters* in the **Methods** section while the default parameters used for all previous experiments can be found in the subsection titled *Transfer learning using DEGAS*. We discovered that among the eight hyper-parameters, the majority of them did not significantly affect the ROC-AUC for predicting GBM subtypes in TCGA GBM patients with the exception of three hyperparameters – namely the drop-out retention rate, number of hidden layer nodes, and *L_2_*-regularization weight with spearman correlation p-value < 0.1 (**Fig. S5, Table S22**). Similarly, the majority of hyper-parameters did not significantly affect the correct assignment of subtype to GBM scRNA-seq tumor, except for a few exceptions in training steps, patient loss weight, and MMD loss weight with spearman correlation p-value < 0.1 (**Fig. S6**, **Table S22**). We therefore suggest users to keep default settings for at least patient loss weight, MMD loss weight, and *L_2_*-regularization weight. The percentage of GBM subtype labels ranking in the top two predicted labels improves from 74% to 82% if the default parameters or greater values are used for patient loss weight, MMD loss weight, and *L_2_*-regularization weight (**Fig. S7**).

### Domain adaptation improves DEGAS disease association transfer

Without any bias, MMD and no MMD performances were not different from one another. After bias was added, MMD did improve the ability of *DEGAS* to transfer disease associations onto cells (**Fig. S8**). MMD is important for our algorithm because the bias added to the patients represents the types of systematic bias that are present between bulk and single cell transcriptomic data. In the example of simulation 2 with cell type 2 bias added, it is clear that all of the patients tended to cluster adjacent to the cell type 2 cluster (**Fig. S8A**). We defined high-risk cells in this example as cells with a disease association >0.2 on a [-1,1] scale. Once the DEGAS model had been trained and the disease associations overlaid onto the cells, the *DEGAS* model trained without MMD predicted many cells in cell types other than cell type 1 as being high-risk (**Fig. S8B**). In contrast, the *DEGAS* model trained with MMD only identified cell type 1 cells opposed to other cell types as high-risk (**Fig. S8C**). Over all 30 experiments, we found that the disease association error was lower in the *DEGAS* models with MMD than in the *DEGAS* models without MMD (t-test p-value < 2.2•10^-16^, **Fig. S8D**). When the cells were ordered by their error, there was no experiment where the *DEGAS* model without MMD consistently outperformed the *DEGAS* model with MMD (Kolmogorov-Smirnov p-value < 2.2•10^-16^, **Fig. S8E**).

### Regularization improves the robustness of DEGAS models

Regularization is an important part of the *DEGAS* model which prevents the data from being overfit. Without regularization, *DEGAS* models perform worse during cross-validation (**Table. S24-25**, **Fig. S9**). There is no case where an unregularized model performed better than a regularized model in predicting patient labels during cross validation. Specifically, in simulation 3, the unregularized models performed 5% worse in PR-AUC when predicting patient labels in patients (**Table. S24**). Similarly, the unregularized *DEGAS* models performed 9% worse in PR-AUC when predicting cell type labels in cells (t-test p-value = 4.34•10^-3^, **Table. S25**). The regularization also improved the transfer of disease associations to the cells in 2/3 simulations (**Fig. S9**).

## Discussion

In this work, we developed the transfer learning framework *DEGAS* to integrate scRNA-seq and patient-level transcriptomic data in order to infer the transferrable “impressions” between patient characteristics in single cells and cellular characteristics in patients. Using transfer learning, we trained a model with both scRNA-seq and patient bulk tissue gene expression data, then reduced the differences between the distributions of the representations for the two data types in the final hidden layer of our model via domain adaptation. This process allows information about patient disease attributes as well as cell types to be transferred between the two data types. We focus on the transfer of patient disease attributes to cells because there are far fewer available methods addressing this task than deconvolution. We tested and validated the *DEGAS* framework on datasets from one simulation and two diseases: GBM, which contained ground truth tumor subtype labels, and AD, which contained ground truth cell type-disease associations.

These experiments on validation datasets demonstrate the necessity for *DEGAS* especially as it relates to the current methods that rely on accurate clustering, cell type annotation, or case-control scRNA-seq. For datasets that contain case and control scRNA-seq data, tools like *Augur* are very effective to prioritize cell types. When no patient level transcriptomic data is available but case-control scRNA-seq is available, tools like *Augur* should be used since *DEGAS* requires patient level transcriptomic data. If patient level transcriptomic data and single cell transcriptomic data are available and there is a necessity to overlay disease associations onto individual cells, then only *DEGAS* can be used. Furthermore, if the scRNA-seq dataset does not contain case and control samples then *DEGAS* needs to be used instead of *Augur* since *Augur* requires case and control samples. The *DEGAS* framework in this sense can be used in a wide variety of study designs as long as there is scRNA-seq and patient transcriptomic data.

Another challenging issue in scRNA-seq analysis is that it is difficult to determine the best clustering options. In our simulation examples, we can determine that the correct number of clusters based on average silhouette width would be four clusters. However, if the number of clusters was increased in the clustering algorithm there would be a stronger correlation between some clusters and disease. Therein lies the challenge – should the clustering results be optimized to reflect the relative transcriptomic signals or should they be optimized to create the greatest correlations with disease state? Furthermore, the different resolutions of clusters may capture different correlations with disease. For these reasons, assigning disease associations directly to cells alleviates some of these problems with cluster resolution decisions. Assigning disease associations directly to cells not only solves the cluster resolution problem but also allows simultaneous identification of cell-intrinsic and cell proportional changes.

The *DEGAS* algorithm can identify both cell-intrinsic changes and cell proportional changes as demonstrated in the simulation examples and the AD study. In simulation 1, the disease is associated with proportional changes in cell type 1. In simulation 2, the disease is associated with a cell-intrinsic change of cell type 4. In simulation 3, there are both cell-intrinsic changes and cell proportional changes in cell type 4. In the AD experiments, two of the single-cell datasets did include data from both AD and normal brains [15, 32]. The cells that came from the AD patients tended to have a higher association with AD, which indicates the detection of cell-intrinsic changes. The importance of cell prioritization at the individual cell level is highlighted in simulation and AD examples. Simulation 2 shows an example where cell level associations are necessary due to clustering results that do not capture the disease associations. Specifically, there are cases where cells will cluster together but have dissimilar associations to disease. If the cells of cell type 4 are not evaluated individually, the association of the cell type 4 disease subtype with disease could be lost. In the AD example, the astrocyte cell type is overall not associated with AD. However, a subset of astrocytes expressing markers for DAAs were found to have a positive disease association while still clustering with the astrocytes that were not associated with disease. Similarly, microglia cells are broadly positively associated with AD but the highest AD association microglia were enriched for DAM markers. When a targeted analysis was performed on only microglia from AD and normal brains, highest AD association microglia were enriched for both HAM and DAM markers. These examples show how DEGAS can identify disease associated cells that cluster within a larger cell type.

In short, the *DEGAS* analysis on AD data further validated our model by correctly identifying the decreased neuron and increased microglia proportions in AD patients. Aside from these known characteristics of AD pathology, we also identified a *GFAP*+ astrocyte subtype taken from normal human brain tissue that is associated with AD and is supported from AD mouse models [55]. We further validated this by finding that *GFAP* expression in Astrocytes was significantly increased in Astrocytes taken from AD patients and concluded that there may be an expansion of this Astrocyte subtype in AD. This is also a convincing example of the utility of *DEGAS* as it assigned disease association at the single cell level, allowing us to identify intra-cell type differences in disease risk that constitute disease-associated cells.

For the GBM single cell patient cohort, each GBM tumor, from which scRNA-seq data was generated, had a GBM subtype label [44]. The *DEGAS* results showed that the majority of cells in each tumor were labeled with the same GBM subtype as previously defined in Patel *et al.* [44]. Specifically, *DEGAS* correctly mapped Proneural, Mesenchymal, Classical, and Neural GBM subtypes to single cells in four GBM tumor samples. This experiment also shows the broad applicability of the model since the single cells had no labels and the patient samples had multiclass labels. *DEGAS* is highly flexible and allows for different categories of output labels to be combined, which may include but are not limited to classification labels, Cox proportional hazard, and even no labels. This allows for a wide variety of applications to adopt the *DEGAS* framework so that impressions are not limited to only one type of disease attribute.

To explore disease with less understood cellular subtypes, we applied *DEGAS* to multiple MM datasets. The models were able to assign PFS metrics to individual cells and subtype populations of CD138+ cells identified by cell type clustering methods Seurat [28] and *BERMUDA* [25]. Among the identified subtypes of cells, subtype 2 was the most consistent between patients visualized by *BERMUDA* (**Fig. S4E**). Furthermore, we found that the subtype 2 cell population appeared to have a gradient of cells moving away from the main subtype 1 group, possibly associated with a certain degree of differentiation (**Fig. S4A-D**). We did experience a lower PR-AUC for subtype 4 than the other subtypes used during model training. However, this subtype was extremely uncommon in the samples and as a result the random PR-AUC would be close to zero making the PR-AUC of 0.44 well above random. Considering that subtype 4 was not found to be highly associated with progression, the lower PR-AUC did not greatly affect our interpretation of the data, which mainly focused on subtype 1 and subtype 2. We believe that *DEGAS* could be improved for highly imbalanced data.

Upon further examination, we found evidence that the subtype 2 cells may represent a population of malignant plasma cells expressing high levels of *PHF19*. *PHF19* is known to play a role in hematopoietic stem cell state and differentiation [65–67] and is a marker for aggressive disease in MM [64]. Furthermore, knock down of *PHF19* has been shown to shift myeloma cells into a less proliferative state [64]. The subtype 2 cells express *SDC1* (also known as *CD138)* and showed significantly increased *PHF19* expression in comparison to the other subtypes. Since all of the IUSM MM cells in our study had already been FACS sorted for CD138+, it is possible we have identified a subpopulation of CD138+/*PHF19^high^* cells in MM tumors. This could prove a useful finding since currently the association between *PHF19* and tumor aggressiveness is at the patient level whereas our results imply that only a fraction of malignant plasma cells in a MM tumor actually overexpress *PHF19*.

This subtype could be targeted using precision immunotherapies that are not restricted to a single patient since the CD138+/*PHF19^high^* cells (*i.e.*, subtype 2) were found to be present in multiple (3/4) patients. Of the three patients with detectable levels of subtype 2 in the CD138+ fraction, two patients (patient 2 and patient 4) had relapsed MM at time of biopsy and the other patient (patient 5) was SMM at biopsy and later progressed to MM. The other patient (patient 3) had little to no detectable subtype 2 cells in the CD138+ fraction and was SMM at time of biopsy and has not progressed to MM. These signs again seem to indicate a common cellular phenotype associated with progression in MM.

Based on the validated results in a variety of disease data analyses, we find that *DEGAS* has broad applications in virtually all diseases with available patient-level and single cell level omic data. The tensorflow [68] machine learning code is integrated with a simple R package interface (https://github.com/tsteelejohnson91/DEGAS) which will facilitate researchers to manipulate scRNA-seq and bulk expression data on their own.

## Conclusion

*DEGAS* is a powerful transfer learning tool for integrating different levels of omic data and identifying the latent molecular relationships between populations of cells and disease attributes, which we refer to as impressions. We validated the *DEGAS* framework on simulated data, GBM and AD by showing *DEGAS* models were capable of accurately predicting patient characteristics at single-cell level. We then leveraged this transfer learning approach on MM data and identified a CD138+/*PHF19^high^* subtype population in MM that was significantly associated with disease progression. This subtype contains unique RNA profiles and gene correlations that could be both leveraged as a prognostic biomarker and possibly targeted directly to reduce the risk of progression. We believe that *DEGAS* can be a powerful solution to overcome the challenge of integrating patient single-cell data with bulk tissue data so that researchers can identify populations of cells associated with an disease attribute of interest. Furthermore, *DEGAS* can accommodate flexible data types. This makes it a highly general framework that can be applied in multiple diseases and data types to identify cellular populations that are associated with prognosis or treatment response, or to identify specific patient groups with certain cell subtypes for personalized treatment.

## Acknowledgements

We thank the Center for Computational Biology and Bioinformatics for the computational resources and work space to complete the research. We also thank the MMRF for the data generated as part of the Multiple Myeloma Research Foundation Personalized Medicine Initiatives (https://research.themmrf.org and www.themmrf.org), the Allen Institute for Brain Science for the data generated as part of their cell types database, and Mount Sinai/JJ Peters VA Medical Center for the data generated as a part of their brain bank.

## Funding

National Institutes of Health NLM-NRSA Fellowship F31LM013056 to TSJ and The Ohio State University (Columbus, OH) and departmental start-up funding from the Indiana University School of Medicine (Indianapolis, IN) to KH and TSJ.

## Author contributions

TSJ, CYY, JZ, and KH conceived and designed the project. TSJ, CYY, SX performed the analyses. TSJ and ZH designed the software package. TSJ, CYY, XH, SX, CD, MA, YW, CB, YZ, YL, JZ, BW, and KH interpreted the results. ZH, TW, WS, YW, and CB provided technical guidance. TSJ, CYY, JZ, and KH wrote the manuscript. JZ and KH supervised the project.

## Competing interests

The authors declare that this research was conducted in the absence of any commercial or financial relationships that could be construed as a potential conflict of interest.

## Data and materials availability

The *DEGAS* R package is freely available on GitHub (https://github.com/tsteelejohnson91/DEGAS). A minimum reproducible example of the IUSM myeloma scRNA-seq data is also deposited on Github. Our complete myeloma scRNA-seq has been deposited on GEO (GSE161722). All other data are publicly available.

## Notes

### Competing Interest Statement

The authors have declared no competing interest.

### Summary of Updates

This version of the manuscript is an updated version which corrects the figure order.

