## Supplementary Materials for "Diagnostic Evidence GAuge of Single cells (DEGAS): A flexible deep-transfer learning framework for prioritizing cells in relation to disease"

### Supplementary Text

#### DEGAS model training algorithm

**Input:** $X_{Pat}$, $X_{Cell}$, $Y_{pat}$, and $Y_{Cell}$, where at least one $Y$ input is present
**Output:** A trained *DEGAS* model ($f_{Class}()$ and/or $f_{Cox}()$, **Eq 2-3**)
Random initialization
Subsample cells and patients (minibatch)
Generated null label cells and patients (see note)
Repeat until convergence:
    Minimize DEGAS loss (**Eq. 8-11**)
    Every 50 iterations:
        Subsample cells and patients (minibatch)
        Generated null label cells and patients (see note)

Note: When cell/patient labels were available, null labelled patients/cells were generated to reinforce the model learning neutral relationships, i.e. non-associations to labels. Given k classes in a multiclass classification problem, k samples were selected using a Gamma distribution based on the labels and used to generate samples with a mixtures of labels. Half of the minibatch would consist of the subsample of cells and the other half would consist of the combination samples derived from them.

#### Disease associated astrocyte (DAA), disease associated microglia (DAM), and human Alzheimer's microglia (HAM) markers

DAA markers were derived from the Habib *et al.* study supplementary table 2 [1]. The markers were included if they were up-regulated in cluster 4 (i.e. DAA cluster). This resulted in 304 DAA markers that were included in the final DAA marker set (**Supplementary File 1**).

HAM markers were derived from the Srinivasan *et. al.* study [2]. The 45 up-regulated markers were used for all of the enrichment analysis of high AD association microglia. For the correlation analysis, both the 45 up-regulated and 23 down-regulated markers were used (**Supplementary File 2**).

DAM markers were derived from the Karen-Shaul *et al.* study Table S2 [3]. The markers were included if they were up-regulated in the DAM cluster and if their -Log10 P was > 10. This resulted in 337 markers (**Supplementary File 3**).

### Supplementary Figures

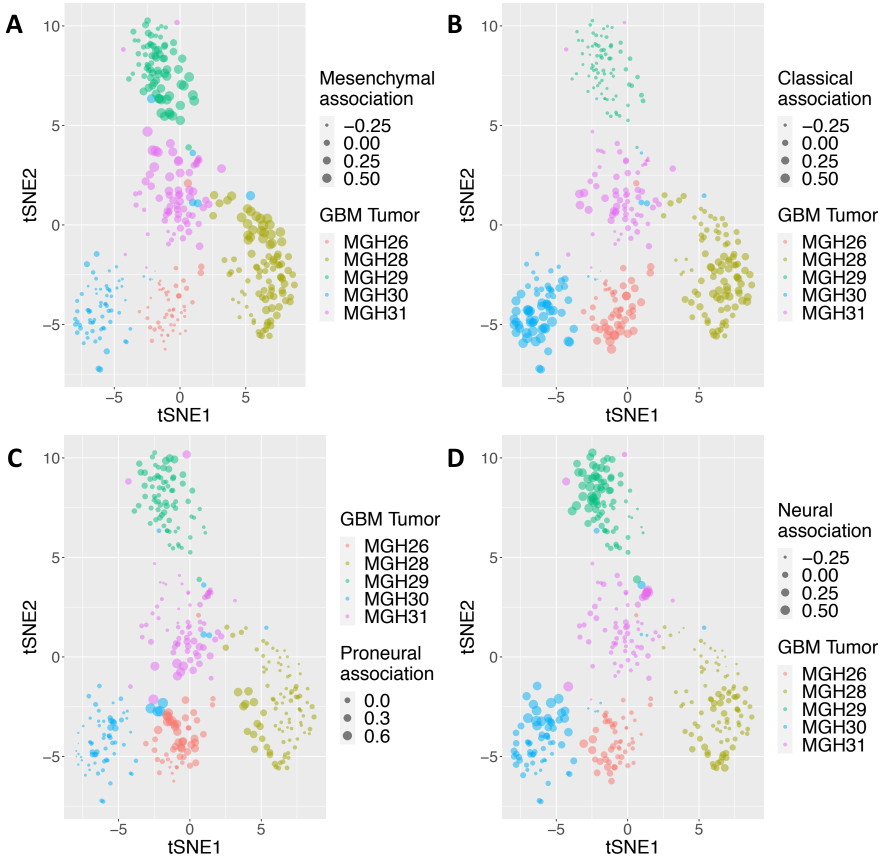


**Fig. S1** Scatterplots for the single cells in each of the Patel et al. GBM tumors overlaid with GBM subtype associations: **A)** Mesenchymal association, **B)** Classical association, **C)** Proneural association, and **D)** Neural association.

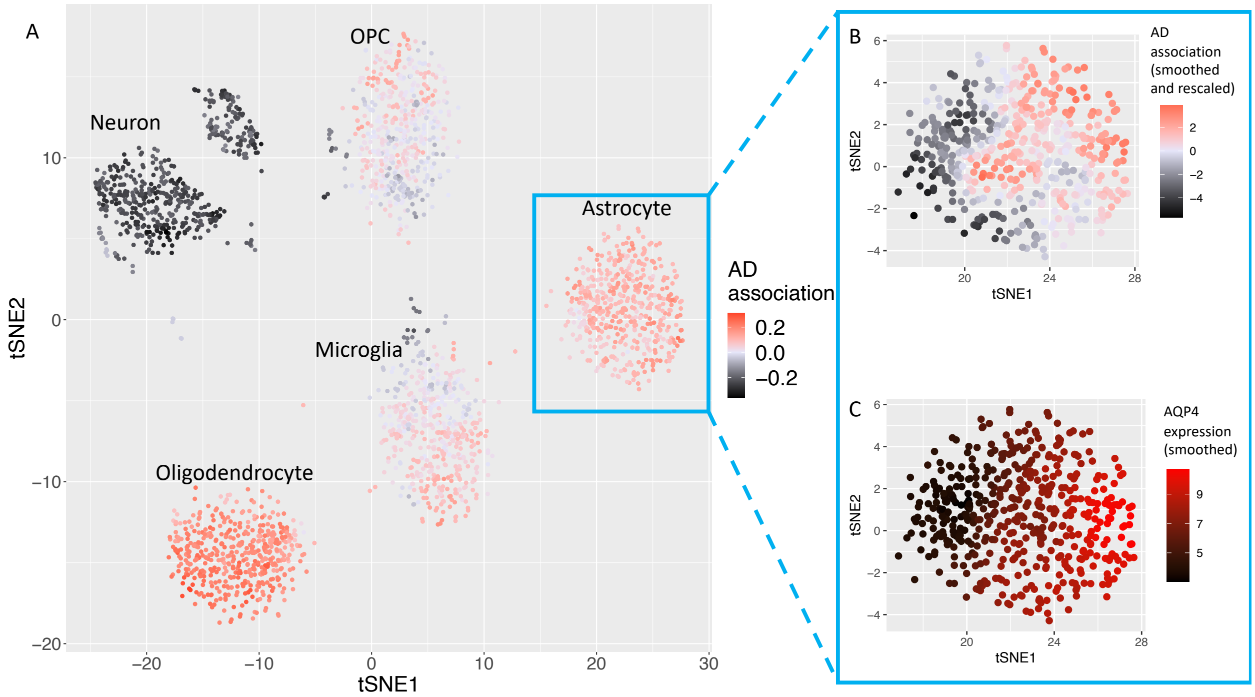


**Fig. S2** Secondary analysis of AIBS single cells. **A)** All cell types overlaid with the AD association scores (sample of 500 per cell type). **B)** AD association overlaid onto only the astrocytes after kNN smoothing. **C)** DAA marker AQP4 overlaid onto the astrocytes after kNN smoothing.

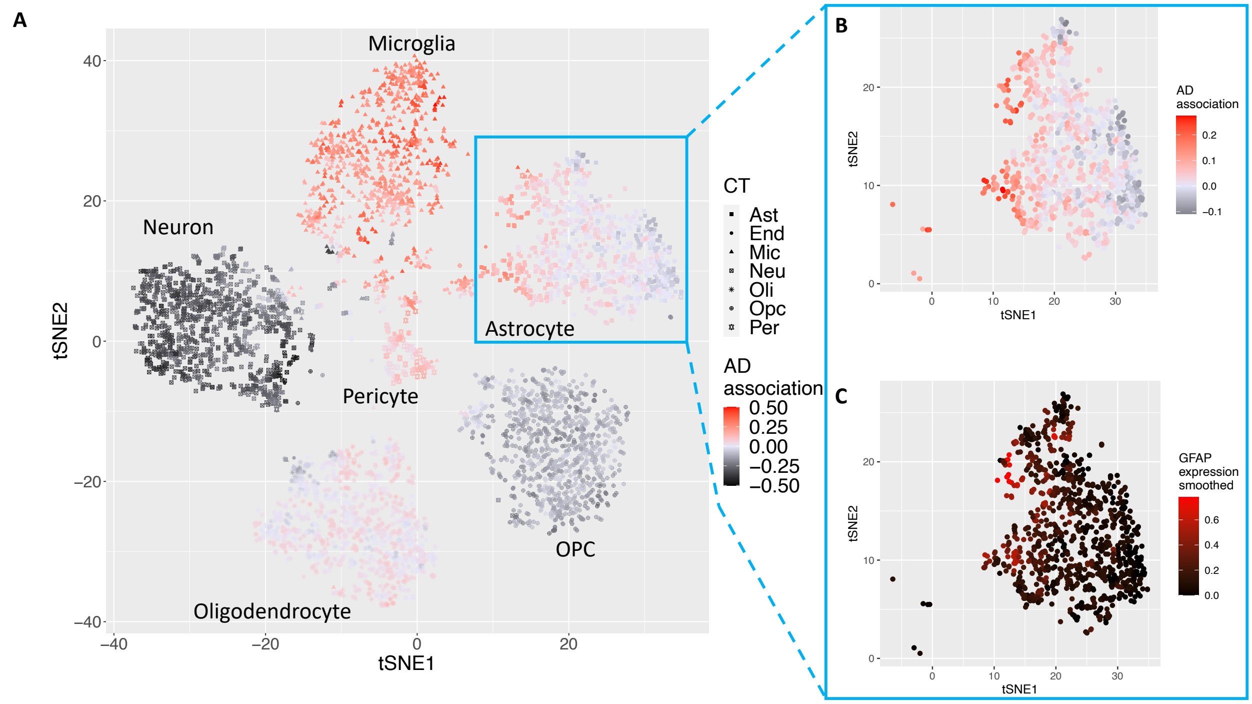


**Fig. S3** Analysis of Mathys et al. single cells. **A)** All cell types overlaid with the AD association scores (sample of 1000 per cell type). **B)** AD association overlaid onto only the astrocytes after kNN smoothing. **C)** DAA marker GFAP overlaid onto the astrocytes after kNN smoothing.

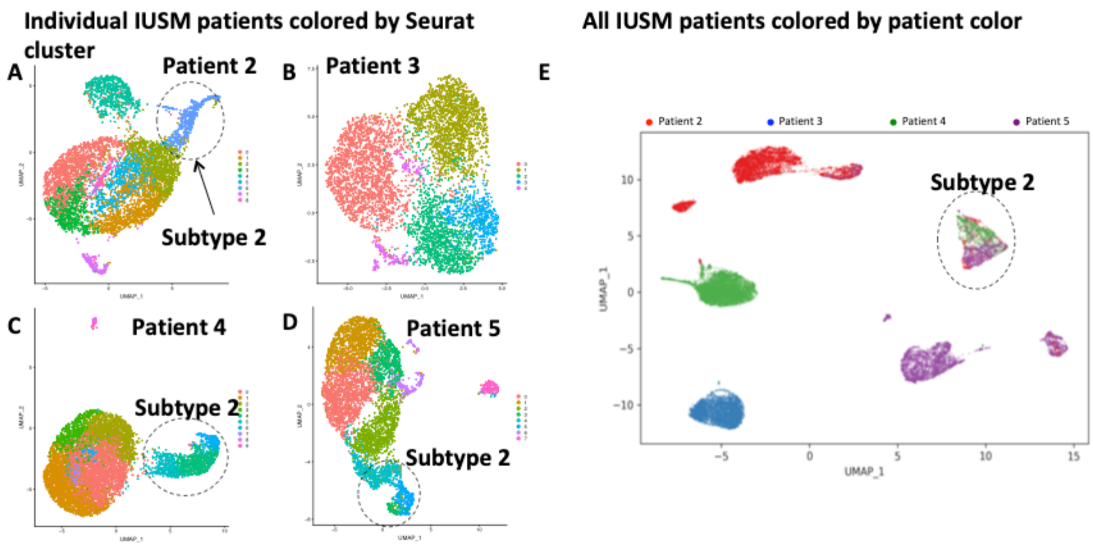


**Fig. S4** Clustering of IUSM myeloma patient single cells. **A-D)** The *Seurat* individual patient clustering results colored by cluster for **A)** IUSM patient 2, **B)** IUSM patient 3, **C)** IUSM patient 4, and **D)** IUSM patient 5. **E)** The clustering results from *BERMUDA* for all four patients colored by patient.

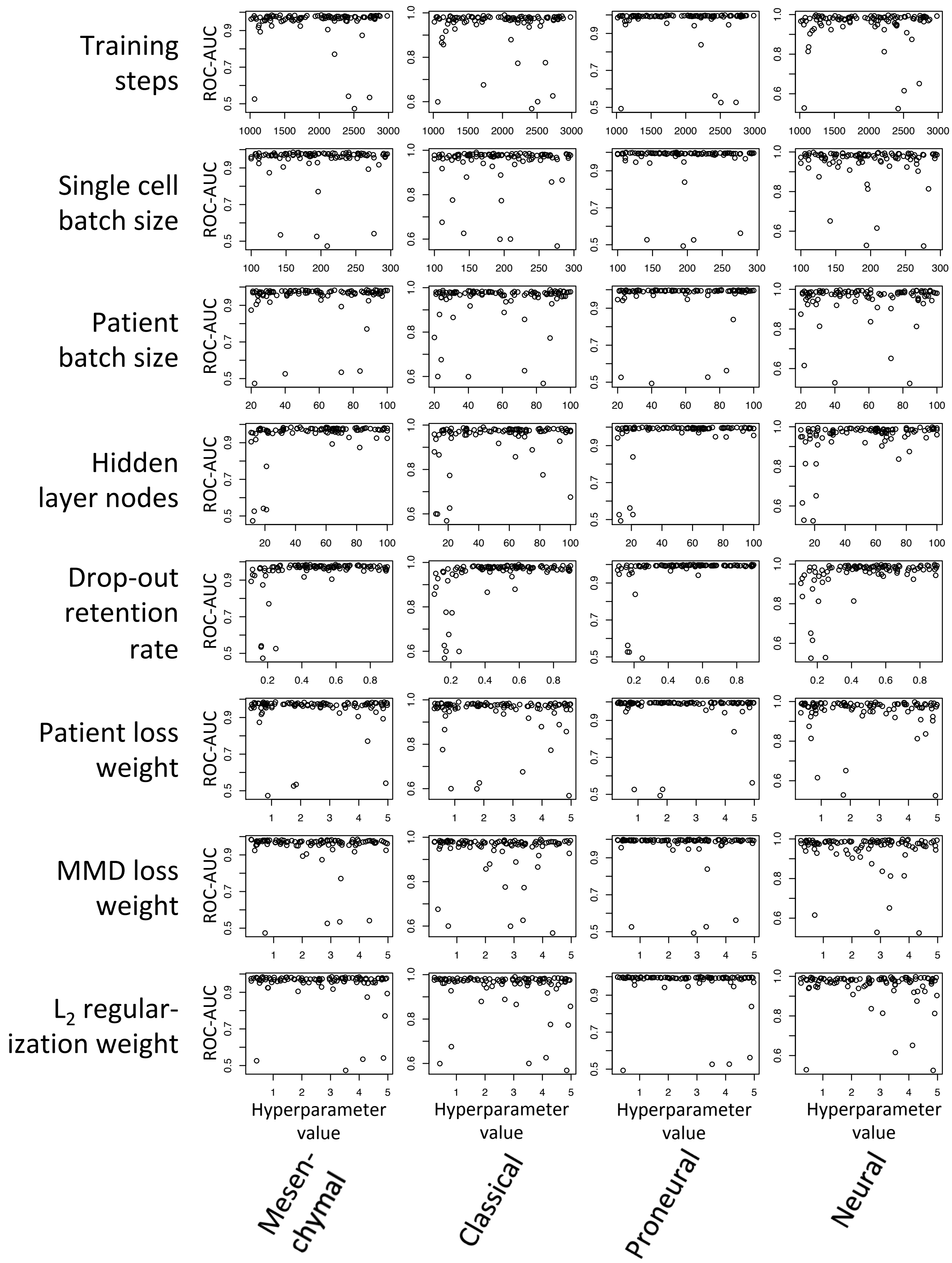


**Fig. S5** All scatterplots between the choices of hyperparameter values and the classification performance. The y-axis is the performance as measured by ROC-AUC for a given GBM subtype label, as shown in the bottom of the figure. Each specific parameter type is shown on the left side of the figure. The corresponding spearman correlation coefficients are computed and shown in **Table S8**.

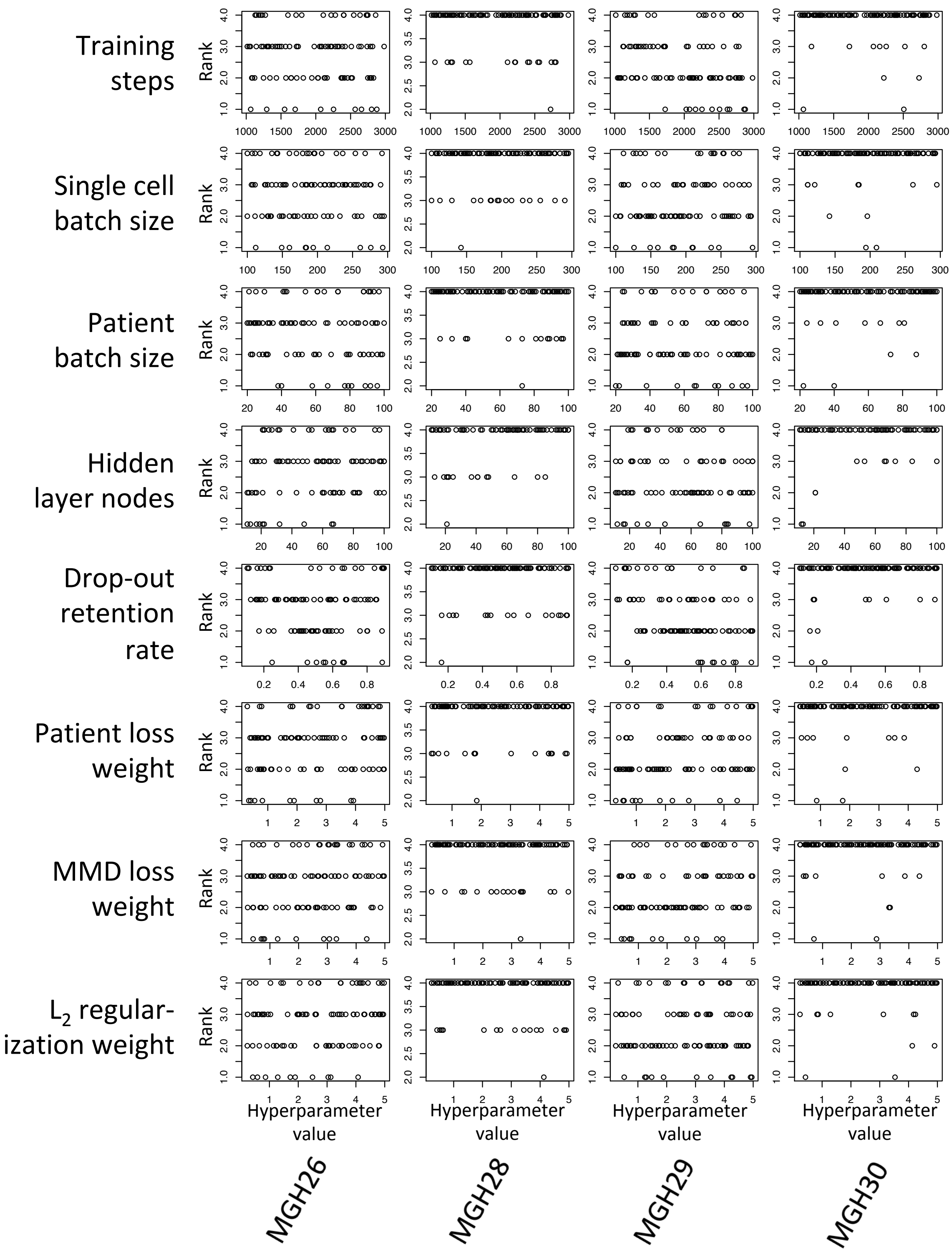


**Fig. S6** All scatterplots between hyperparameter values and correct label rank such that 4 is the desired rank. The x-axis values in each plot is the hyperparameter value of a specific hyperparameter label, as shown on the left side of the figure. The y-axis is the rank for the correct label in a given GBM tumor. The corresponding spearman correlation coefficients are computed and shown in **Table S9**.

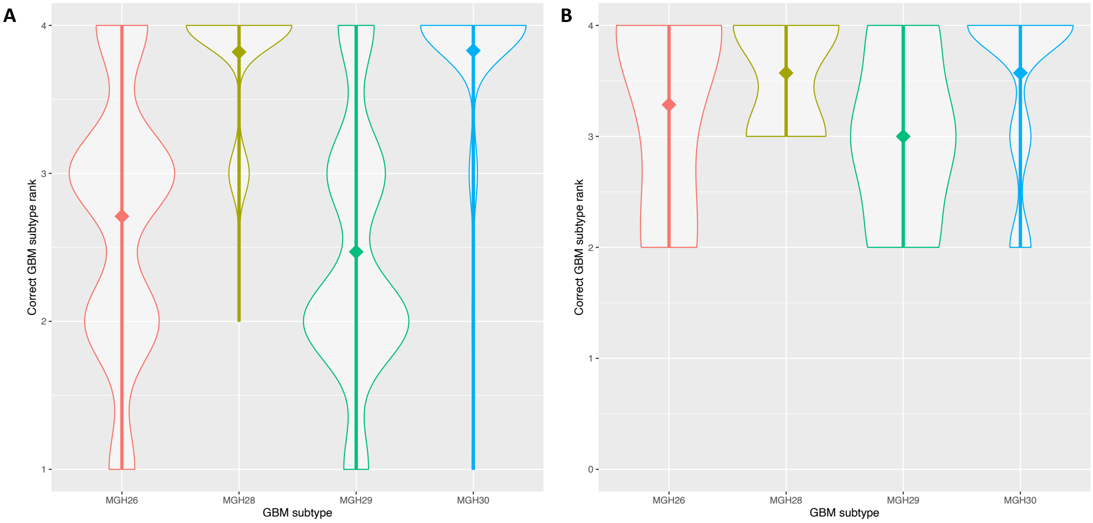


**Fig. S7** Rank of the correct cell label for each of the Patel et al. GBM tumors. **A)** These are the results including all 100 randomly selected sets of hyperparameters. **B)** These are the results including only the hyperparameters where $\lambda_1$, $\lambda_2$, and $\lambda_3$ are greater than 3 that represent the default hyperparameters. The rank of the correct label was calculated by calculating the mean of each GBM subtype association across all of the cells in that tumor. This resulted in each of the 100 random hyperparameters having a rank for each GBM subtype for each of the GBM scRNA-seq tumors (4 highest ranked, 1 lowest ranked). Ideally the rank for correct labels would be 4 since there are 4 total labels (Mesenchymal, Classical, Proneural, Neural). The mean rank value is marked by a diamond in both **A** and **B**.

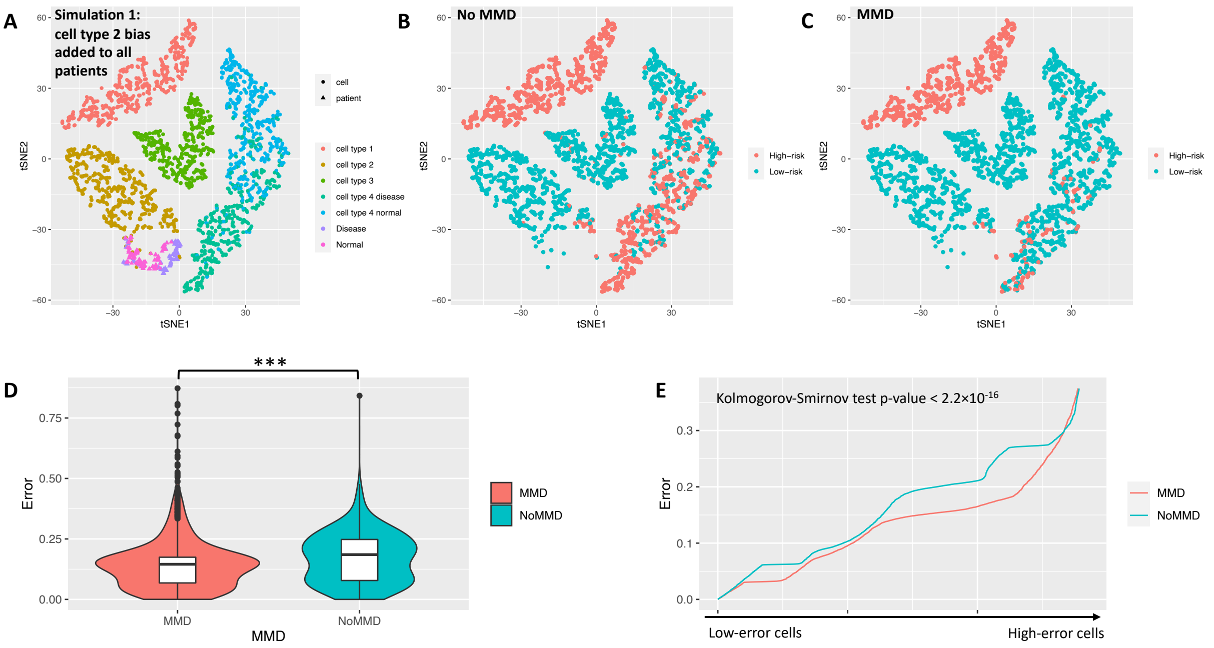


**Fig. S8** Necessity of domain adaptation to transfer disease associations from patients to cells. **A)** The simulation 1 cells and patients with cell type 2 systematic bias added to all of the patients. The disease associations stratified into high-risk and low-risk cells based on a 0.2 cutoff from **B)** the model without domain adaptation (No MMD) and **C)** the model with domain adaptation (MMD). **D)** The individual cell errors across the three simulations comparing models without domain adaptation (No MMD) and with domain adaptation (MMD). **E)** The cells sorted by error to show that there is little crossover between the two model errors. *** denotes p-value <0.001

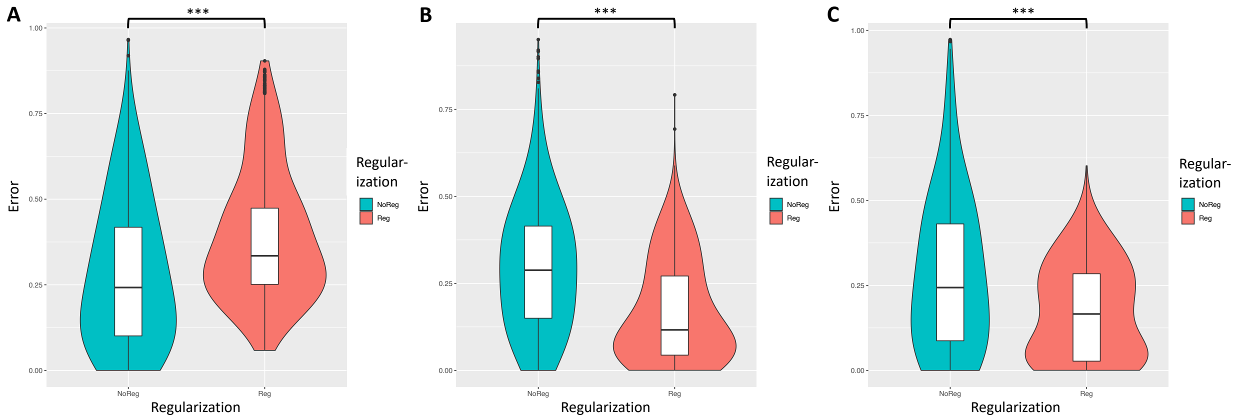


**Fig. S9** The necessity of regularization to transfer disease associations from patients to cells. Individual cell errors from cross validation comparing models with and without regularization in **A)** simulation 1, **B)** simulation 2, and **C)** simulation 3. *** denotes p-value <0.001

**Supplementary Tables**

**Table S1.** Prediction performance metrics for simulated patients including receiver operating curve (ROC) area under the curve (AUC) and precision recall (PR) curve AUC.

| | **Simulation 1** | | **Simulation 2** | | **Simulation 3** | |
|---|---|---|---|---|---|---|
| | **ROC-AUC** | **PR-AUC** | **ROC-AUC** | **PR-AUC** | **ROC-AUC** | **PR-AUC** |
| **Disease status** | 0.94 | 0.96 | 0.99 | 0.99 | 0.96 | 0.98 |

**Table S2.** Prediction performance metrics for simulated cells including ROC-AUC and PR-AUC.

| | **Simulation 1** | | **Simulation 2** | | **Simulation 3** | |
|---|---|---|---|---|---|---|
| | **ROC-AUC** | **PR-AUC** | **ROC-AUC** | **PR-AUC** | **ROC-AUC** | **PR-AUC** |
| **Cell type 1** | 1.00 | 1.00 | 1.00 | 1.00 | 1.00 | 1.00 |
| **Cell type 2** | 1.00 | 1.00 | 1.00 | 1.00 | 1.00 | 1.00 |
| **Cell type 3** | 1.00 | 1.00 | 1.00 | 1.00 | 1.00 | 1.00 |
| **Cell type 4** | 1.00 | 1.00 | 1.00 | 1.00 | 1.00 | 1.00 |

**Table S3.** Prediction performance metrics for TCGA GBM patients into molecular subtypes including ROC-AUC and PR-AUC.

| | **ROC-AUC** | **PR-AUC** |
|---|---|---|
| **Mesenchymal** | 0.93 | 0.89 |
| **Classical** | 0.94 | 0.86 |
| **Proneural** | 0.99 | 0.97 |
| **Neural** | 0.97 | 0.79 |

**Table S4.** Prediction performance metrics for MSBB patients including ROC-AUC and PR-AUC for both the primary analysis (Seurat feature selection) and secondary analysis (sparsity- and variance-based feature selection.

| | **Primary Analysis** | | **Secondary Analysis** | |
|---|---|---|---|---|
| | **ROC-AUC** | **PR-AUC** | **ROC-AUC** | **PR-AUC** |
| **AD status** | 0.78 | 0.82 | 0.68 | 0.76 |

**Table S5.** Prediction performance metrics for AIBS cells including ROC-AUC and PR-AUC for both the primary analysis (Seurat feature selection) and secondary analysis (sparsity- and variance-based feature selection.

| | **Primary Analysis** | | **Secondary Analysis** | |
|---|---|---|---|---|
| | **ROC-AUC** | **PR-AUC** | **ROC-AUC** | **PR-AUC** |
| **Neuron** | 0.99 | 0.99 | 0.99 | 0.99 |
| **Oligodendrocyte** | 0.99 | 0.99 | 0.99 | 0.99 |
| **Astrocyte** | 0.99 | 0.99 | 0.99 | 0.99 |
| **OPC** | 0.99 | 0.99 | 0.99 | 0.99 |
| **Microglia** | 0.99 | 0.99 | 0.99 | 0.99 |

**Table S6.** Significant differentially expressed DAA markers in primary analysis of AIBS Institute astrocytes for high AD association ($AD^{high}$) vs. low AD association ($AD^{low}$) astrocyte cells.

| | **$AD^{high}$ expressed proportion** | **$AD^{low}$ expressed proportion** | **FC** | **t-test p-value** |
|---|---|---|---|---|
| **GFAP** | 0.74 | 0.48 | 2.52 | $1.78 \cdot 10^{-13}$ |
| **CRYAB** | 0.45 | 0.26 | 4.07 | $6.83 \cdot 10^{-11}$ |

| | | | | |
|---|---|---|---|---|
| **S100B** | 0.71 | 0.56 | 1.15 | $6.20 \cdot 10^{-9}$ |
| **DBI** | 0.54 | 0.37 | 1.56 | $9.09 \cdot 10^{-7}$ |
| **FOS** | 0.27 | 0.11 | 3.54 | $1.16 \cdot 10^{-6}$ |
| **JUND** | 0.30 | 0.14 | 2.09 | $1.77 \cdot 10^{-6}$ |
| **ID2** | 0.76 | 0.62 | 0.76 | $2.05 \cdot 10^{-6}$ |
| **CST3** | 0.95 | 0.91 | 0.46 | $1.37 \cdot 10^{-5}$ |
| **JUNB** | 0.22 | 0.09 | 3.08 | $2.02 \cdot 10^{-5}$ |
| **IGFBP5** | 0.17 | 0.05 | 6.47 | $2.41 \cdot 10^{-5}$ |
| **ITM2B** | 0.61 | 0.46 | 0.79 | $2.78 \cdot 10^{-5}$ |
| **IFITM3** | 0.44 | 0.25 | 1.03 | $6.42 \cdot 10^{-5}$ |
| **B2M** | 0.36 | 0.28 | 1.41 | $1.11 \cdot 10^{-4}$ |
| **ID4** | 0.63 | 0.53 | 0.55 | $2.50 \cdot 10^{-4}$ |
| **PRDX1** | 0.40 | 0.24 | 0.84 | $5.35 \cdot 10^{-4}$ |
| **RHOB** | 0.40 | 0.27 | 0.93 | $1.14 \cdot 10^{-3}$ |
| **PPIB** | 0.17 | 0.10 | 1.18 | $1.18 \cdot 10^{-3}$ |
| **SLC38A1** | 0.25 | 0.09 | 1.35 | $1.20 \cdot 10^{-3}$ |
| **AGT** | 0.62 | 0.47 | 0.50 | $1.59 \cdot 10^{-3}$ |
| **SCG3** | 0.64 | 0.59 | 0.55 | $1.83 \cdot 10^{-3}$ |
| **ID3** | 0.24 | 0.09 | 1.39 | $1.99 \cdot 10^{-3}$ |
| **SPARC** | 0.16 | 0.06 | 2.91 | $2.06 \cdot 10^{-3}$ |
| **SOX2** | 0.81 | 0.82 | 0.35 | $2.80 \cdot 10^{-3}$ |
| **ARHGEF4** | 0.70 | 0.61 | 0.43 | $3.61 \cdot 10^{-3}$ |
| **FBXO2** | 0.24 | 0.15 | 0.93 | $3.64 \cdot 10^{-3}$ |
| **CLU** | 0.97 | 0.94 | 0.37 | $4.24 \cdot 10^{-3}$ |
| **SCRG1** | 0.36 | 0.29 | 0.73 | $4.68 \cdot 10^{-3}$ |
| **NPC2** | 0.13 | 0.06 | 1.55 | $6.30 \cdot 10^{-3}$ |
| **AQP4** | 0.85 | 0.70 | 0.38 | $6.35 \cdot 10^{-3}$ |
| **PRDX6** | 0.43 | 0.39 | 0.48 | $8.05 \cdot 10^{-3}$ |
| **APOE** | 0.71 | 0.61 | 0.45 | $9.04 \cdot 10^{-3}$ |
| **LGALS1** | 0.13 | 0.07 | 1.30 | $9.33 \cdot 10^{-3}$ |
| **FABP7** | 0.26 | 0.16 | 0.77 | $1.32 \cdot 10^{-2}$ |
| **CPE** | 0.82 | 0.69 | 0.33 | $1.40 \cdot 10^{-2}$ |
| **STAT3** | 0.20 | 0.17 | 0.86 | $1.52 \cdot 10^{-2}$ |
| **TUBB2B** | 0.39 | 0.34 | 0.44 | $1.88 \cdot 10^{-2}$ |
| **CD81** | 0.48 | 0.42 | 0.40 | $2.44 \cdot 10^{-2}$ |
| **HRSP12** | 0.19 | 0.12 | 0.91 | $2.57 \cdot 10^{-2}$ |
| **TMBIM6** | 0.42 | 0.31 | 0.50 | $2.76 \cdot 10^{-2}$ |
| **UBC** | 0.35 | 0.22 | 0.76 | $2.93 \cdot 10^{-2}$ |
| **AEBP1** | 0.34 | 0.25 | 0.51 | $3.23 \cdot 10^{-2}$ |

| | | | | |
|---|---|---|---|---|
| **VIM** | 0.29 | 0.19 | 0.59 | $3.37 \bullet 10^{-2}$ |
| **SOX9** | 0.49 | 0.45 | 0.36 | $3.47 \bullet 10^{-2}$ |
| **GSN** | 0.45 | 0.31 | 0.42 | $4.92 \bullet 10^{-2}$ |

**Table S7**. Significant differentially expressed DAM markers in the primary analysis of AIBS microglia for high AD association ($AD^{high}$) vs. low AD association ($AD^{low}$) microglia cells.

| | **$AD^{high}$ expressed proportion** | **$AD^{low}$ expressed proportion** | **FC** | **t-test p-value** |
|---|---|---|---|---|
| **CD74** | 0.91 | 0.92 | 0.47 | $2.21 \bullet 10^{-5}$ |
| **RPL10A** | 0.15 | 0.12 | 1.72 | $1.46 \bullet 10^{-3}$ |
| **GM2A** | 0.11 | 0.06 | 3.63 | $1.48 \bullet 10^{-3}$ |
| **SPP1** | 0.59 | 0.50 | 0.68 | $1.71 \bullet 10^{-3}$ |
| **B2M** | 0.70 | 0.61 | 0.42 | $5.33 \bullet 10^{-3}$ |
| **RPL37** | 0.17 | 0.13 | 1.27 | $6.83 \bullet 10^{-3}$ |
| **CXCL16** | 0.19 | 0.12 | 1.02 | $1.77 \bullet 10^{-2}$ |
| **TREM2** | 0.39 | 0.26 | 0.46 | $3.39 \bullet 10^{-2}$ |
| **RPS14** | 0.26 | 0.20 | 0.85 | $3.41 \bullet 10^{-2}$ |
| **APOE** | 0.37 | 0.32 | 0.42 | $3.83 \bullet 10^{-2}$ |
| **RPL7** | 0.12 | 0.11 | 0.85 | $4.57 \bullet 10^{-2}$ |

**Table S8.** Significant differentially expressed DAA markers in secondary analysis of AIBS astrocytes for high AD association ($AD^{high}$) vs. low AD association ($AD^{low}$) astrocyte cells.

| | **$AD^{high}$ expressed proportion** | **$AD^{low}$ expressed proportion** | **FC** | **t-test p-value** |
|---|---|---|---|---|
| **ADD3** | 0.86 | 0.80 | 0.29 | $2.82 \bullet 10^{-4}$ |
| **HSD17B4** | 0.34 | 0.27 | 0.61 | $1.80 \bullet 10^{-3}$ |
| **MFAP3L** | 0.52 | 0.46 | 0.56 | $2.35 \bullet 10^{-3}$ |
| **MID1** | 0.29 | 0.19 | 0.75 | $2.68 \bullet 10^{-3}$ |
| **CRYAB** | 0.48 | 0.34 | 0.55 | $5.46 \bullet 10^{-3}$ |
| **CLDND1** | 0.23 | 0.15 | 1.28 | $7.43 \bullet 10^{-3}$ |
| **LHFP** | 0.12 | 0.10 | 0.95 | $7.44 \bullet 10^{-3}$ |
| **ABCA1** | 0.38 | 0.34 | 0.53 | $1.12 \bullet 10^{-2}$ |
| **USP24** | 0.38 | 0.29 | 0.51 | $1.22 \bullet 10^{-2}$ |
| **TST** | 0.10 | 0.06 | 1.11 | $1.50 \bullet 10^{-2}$ |
| **RASSF8** | 0.18 | 0.12 | 0.79 | $2.34 \bullet 10^{-2}$ |
| **VIMP** | 0.20 | 0.14 | 0.54 | $2.82 \bullet 10^{-2}$ |
| **S100A16** | 0.18 | 0.13 | 0.51 | $2.93 \bullet 10^{-2}$ |
| **HSPA2** | 0.10 | 0.08 | 1.00 | $3.16 \bullet 10^{-2}$ |
| **CYR61** | 0.11 | 0.08 | 0.82 | $3.72 \bullet 10^{-2}$ |

| | | | | |
|---|---|---|---|---|
| **RORB** | 0.25 | 0.19 | 0.52 | $4.02 \cdot 10^{-2}$ |
| **EFR3B** | 0.44 | 0.38 | 0.31 | $4.05 \cdot 10^{-2}$ |
| **TTC8** | 0.11 | 0.09 | 0.89 | $4.46 \cdot 10^{-2}$ |

**Table S9**. Significant differentially expressed DAM markers in the secondary analysis of AIBS microglia for high AD association ($AD^{high}$) vs. low AD association ($AD^{low}$) microglia cells.

| | **$AD^{high}$ expressed proportion** | **$AD^{low}$ expressed proportion** | **FC** | **t-test p-value** |
|---|---|---|---|---|
| **B2M** | 0.79 | 0.52 | 1.57 | $1.06 \cdot 10^{-12}$ |
| **CD74** | 0.92 | 0.90 | 0.59 | $2.85 \cdot 10^{-7}$ |
| **EEF1A1** | 0.40 | 0.34 | 1.04 | $2.52 \cdot 10^{-4}$ |
| **NPC2** | 0.18 | 0.10 | 1.52 | $1.24 \cdot 10^{-3}$ |
| **GRN** | 0.31 | 0.21 | 1.10 | $1.80 \cdot 10^{-3}$ |
| **CSF2RA** | 0.54 | 0.51 | 0.57 | $6.68 \cdot 10^{-3}$ |
| **CTSL** | 0.13 | 0.10 | 1.28 | $1.12 \cdot 10^{-2}$ |
| **TYROBP** | 0.50 | 0.42 | 0.41 | $1.35 \cdot 10^{-2}$ |
| **GPR65** | 0.14 | 0.10 | 2.02 | $1.40 \cdot 10^{-2}$ |
| **CTSB** | 0.72 | 0.66 | 0.35 | $2.60 \cdot 10^{-2}$ |
| **RPL10A** | 0.16 | 0.13 | 0.91 | $2.86 \cdot 10^{-2}$ |
| **TREM2** | 0.37 | 0.29 | 0.42 | $4.67 \cdot 10^{-2}$ |
| **CLEC7A** | 0.45 | 0.32 | 0.42 | $4.94 \cdot 10^{-2}$ |

**Table S10.** Significant differentially expressed DAA markers in Grubman *et al.* astrocytes for high AD association ($AD^{high}$) vs. low AD association ($AD^{low}$) astrocyte cells.

| | **$AD^{high}$ expressed proportion** | **$AD^{low}$ expressed proportion** | **FC** | **t-test p-value** |
|---|---|---|---|---|
| **GFAP** | 0.73 | 0.30 | 4.54 | $2.02 \cdot 10^{-101}$ |
| **CRYAB** | 0.57 | 0.31 | 2.49 | $8.16 \cdot 10^{-49}$ |
| **SLC38A1** | 0.21 | 0.03 | 9.59 | $2.41 \cdot 10^{-29}$ |
| **PLCE1** | 0.24 | 0.08 | 2.69 | $1.77 \cdot 10^{-23}$ |
| **FOS** | 0.33 | 0.15 | 1.62 | $1.38 \cdot 10^{-20}$ |
| **AEBP1** | 0.28 | 0.14 | 1.60 | $2.18 \cdot 10^{-20}$ |
| **IFITM3** | 0.27 | 0.13 | 1.59 | $5.63 \cdot 10^{-19}$ |
| **ID3** | 0.27 | 0.08 | 2.57 | $4.04 \cdot 10^{-17}$ |
| **SORBS1** | 0.83 | 0.76 | 0.38 | $1.54 \cdot 10^{-16}$ |
| **ITPKB** | 0.47 | 0.35 | 0.66 | $8.26 \cdot 10^{-15}$ |
| **ARHGEF4** | 0.58 | 0.48 | 0.52 | $2.77 \cdot 10^{-14}$ |
| **UBC** | 0.50 | 0.36 | 0.81 | $3.03 \cdot 10^{-14}$ |
| **JUNB** | 0.27 | 0.17 | 0.98 | $1.57 \cdot 10^{-7}$ |

**Table S11.** Significant differentially expressed DAM markers in Grubman *et al.* microglia for high AD association ($AD^{high}$) vs. low AD association ($AD^{low}$) microglia cells.

| | $AD^{high}$ expressed proportion | $AD^{low}$ expressed proportion | FC | t-test p-value |
|---|---|---|---|---|
| **SPP1** | 0.60 | 0.41 | 2.65 | $1.46•10^{-7}$ |
| **CSF2RA** | 0.42 | 0.34 | 0.48 | $7.18•10^{-3}$ |
| **HIF1A** | 0.25 | 0.20 | 0.66 | $2.19•10^{-2}$ |

**Table S12.** Significant correlations of kNN smoothed HAM markers with AD association score in targeted analysis of Grubman *et al.* microglia.

| | HAM | PCC | PCC p-value |
|---|---|---|---|
| **CBX6** | HAM up-regulated | 0.11 | $2.34•10^{-2}$ |
| **CLDN15** | HAM up-regulated | 0.14 | $3.29•10^{-3}$ |
| **TSHZ3** | HAM up-regulated | 0.15 | $1.34•10^{-3}$ |
| **DPYD** | HAM up-regulated | 0.39 | $4.46•10^{-18}$ |
| **PTPRG** | HAM up-regulated | 0.31 | $2.63•10^{-11}$ |
| **ACD** | HAM up-regulated | -0.17 | $1.98•10^{-4}$ |
| **RFX2** | HAM up-regulated | 0.20 | $1.42•10^{-5}$ |
| **APOE** | HAM up-regulated | 0.18 | $1.15•10^{-4}$ |
| **ATOH8** | HAM up-regulated | 0.16 | $9.72•10^{-4}$ |
| **GAS2L1** | HAM up-regulated | 0.19 | $3.22•10^{-5}$ |
| **PTPRZ1** | HAM down-regulated | -0.22 | $3.77•10^{-6}$ |
| **SELENBP1** | HAM down-regulated | -0.13 | $7.79•10^{-3}$ |

**Table S13.** Significant differentially expressed HAM markers in targeted analysis of Grubman *et al.* microglia for high AD association ($AD^{high}$) vs. low AD association ($AD^{low}$) microglia cells.

| | $AD^{high}$ expressed proportion | $AD^{low}$ expressed proportion | FC | t-test p-value |
|---|---|---|---|---|
| **PTPRG** | 0.16 | 0.06 | 3.07 | $6.14•10^{-4}$ |
| **APOE** | 0.60 | 0.52 | 0.43 | $2.29•10^{-2}$ |
| **DPYD** | 0.37 | 0.28 | 0.45 | $4.57•10^{-2}$ |

**Table S14.** Significant differentially expressed DAM markers in targeted analysis of Grubman *et al.* microglia for high AD association ($AD^{high}$) vs. low AD association ($AD^{low}$) microglia cells.

| | $AD^{high}$ expressed proportion | $AD^{low}$ expressed proportion | FC | t-test p-value |
|---|---|---|---|---|
| **SPP1** | 0.67 | 0.34 | 5.12 | $3.54•10^{-11}$ |
| **RPL6** | 0.21 | 0.10 | 1.89 | $1.63•10^{-4}$ |
| **RPS28** | 0.28 | 0.14 | 1.28 | $1.65•10^{-4}$ |
| **RPS4X** | 0.22 | 0.11 | 1.36 | $9.15•10^{-4}$ |
| **MYO1E** | 0.21 | 0.09 | 1.57 | $1.45•10^{-3}$ |

| | | | | |
|---|---|---|---|---|
| **RPL32** | 0.25 | 0.16 | 0.99 | $4.95 \cdot 10^{-3}$ |
| **TPT1** | 0.21 | 0.14 | 1.01 | $5.75 \cdot 10^{-3}$ |
| **RPS20** | 0.21 | 0.12 | 1.18 | $6.33 \cdot 10^{-3}$ |
| **FTH1** | 0.61 | 0.58 | 0.70 | $9.02 \cdot 10^{-3}$ |
| **RPS16** | 0.21 | 0.12 | 0.82 | $1.25 \cdot 10^{-2}$ |
| **RPLP2** | 0.18 | 0.12 | 0.89 | $1.33 \cdot 10^{-2}$ |
| **APOE** | 0.60 | 0.52 | 0.43 | $2.29 \cdot 10^{-2}$ |
| **RPL27A** | 0.20 | 0.13 | 0.85 | $2.61 \cdot 10^{-2}$ |
| **RPL37** | 0.13 | 0.08 | 1.01 | $2.90 \cdot 10^{-2}$ |
| **TYROBP** | 0.25 | 0.16 | 0.65 | $3.06 \cdot 10^{-2}$ |
| **RPL7** | 0.19 | 0.15 | 0.69 | $3.42 \cdot 10^{-2}$ |

**Table S15.** Prediction performance metrics for MSBB patients including ROC-AUC and PR-AUC in the Mathys *et al.* analysis.

| | **ROC-AUC** | **PR-AUC** |
|---|---|---|
| **AD status** | 0.77 | 0.81 |

**Table S16.** Prediction performance metrics for Mathys *et al.* cells including ROC-AUC and PR-AUC.

| | **ROC-AUC** | **PR-AUC** |
|---|---|---|
| **Neuron** | 0.99 | 0.99 |
| **Oligodendrocyte** | 0.99 | 0.99 |
| **Astrocyte** | 0.99 | 0.99 |
| **OPC** | 0.99 | 0.99 |
| **Microglia** | 0.99 | 0.99 |
| **Endothelial** | 0.99 | 0.82 |
| **Pericyte** | 0.99 | 0.91 |

**Table S17.** Comparison of AD association scores with AD diagnosis and diagnostic scores for each cell type in the Mathys analysis.

| | | **AD status (AD vs. Normal cells)** | | | **AD association vs. neuritic plaque number** | |
|---|---|---|---|---|---|---|
| **Cell type** | **Number of cells** | **Mean in AD cells** | **Mean in Normal cells** | **t-test p-value** | **PCC** | **PCC p-value** |
| Neu | 1000 | -0.28 | -0.30 | $2.79 \cdot 10^{-6}$ | 0.11 | $3.20 \cdot 10^{-4}$ |
| Oli | 1000 | 0.03 | 0.02 | $2.37 \cdot 10^{-4}$ | 0.09 | $2.75 \cdot 10^{-3}$ |
| Ast | 1000 | 0.04 | 0.03 | $1.09 \cdot 10^{-3}$ | 0.22 | $1.14 \cdot 10^{-2}$ |
| Opc | 1000 | -0.11 | -0.12 | $9.18 \cdot 10^{-4}$ | 0.12 | $1.98 \cdot 10^{-4}$ |
| Mic | 1000 | 0.25 | 0.22 | $2.47 \cdot 10^{-4}$ | 0.13 | $1.89 \cdot 10^{-5}$ |
| End | 121 | 0.16 | 0.09 | $6.33 \cdot 10^{-4}$ | 0.28 | $2.13 \cdot 10^{-3}$ |
| Per | 167 | 0.06 | 0.06 | $7.99 \cdot 10^{-1}$ | -0.02 | $7.70 \cdot 10^{-1}$ |

**Table S18.** Significant differentially expressed DAA markers in the Mathys et al. astrocytes for high AD association ($AD^{high}$) vs. low AD association ($AD^{low}$) astrocyte cells.

| | $AD^{high}$ expressed proportion | $AD^{low}$ expressed proportion | FC | t-test p-value |
|---|---|---|---|---|
| **GFAP** | 0.60 | 0.45 | 0.81 | $1.40 \cdot 10^{-10}$ |
| **CRYAB** | 0.29 | 0.16 | 1.74 | $2.46 \cdot 10^{-10}$ |
| **CLU** | 0.83 | 0.74 | 0.55 | $1.49 \cdot 10^{-9}$ |
| **AQP4** | 0.73 | 0.63 | 0.43 | $3.67 \cdot 10^{-7}$ |
| **UBC** | 0.31 | 0.23 | 0.74 | $1.29 \cdot 10^{-5}$ |
| **PRDX6** | 0.25 | 0.16 | 0.69 | $2.44 \cdot 10^{-4}$ |
| **ITPKB** | 0.61 | 0.52 | 0.29 | $9.49 \cdot 10^{-4}$ |
| **S100A6** | 0.11 | 0.06 | 1.27 | $1.81 \cdot 10^{-3}$ |
| **PLCE1** | 0.15 | 0.09 | 0.85 | $2.31 \cdot 10^{-3}$ |
| **VIM** | 0.26 | 0.19 | 0.52 | $2.98 \cdot 10^{-3}$ |
| **FBXO2** | 0.17 | 0.13 | 0.64 | $8.41 \cdot 10^{-3}$ |
| **ID3** | 0.10 | 0.05 | 1.03 | $1.11 \cdot 10^{-2}$ |
| **MLC1** | 0.40 | 0.34 | 0.26 | $1.64 \cdot 10^{-2}$ |
| **AEBP1** | 0.20 | 0.15 | 0.43 | $2.25 \cdot 10^{-2}$ |
| **GLIS3** | 0.71 | 0.64 | 0.16 | $2.28 \cdot 10^{-2}$ |
| **DBI** | 0.20 | 0.16 | 0.39 | $3.02 \cdot 10^{-2}$ |
| **ID2** | 0.34 | 0.27 | 0.32 | $3.29 \cdot 10^{-2}$ |
| **S100B** | 0.23 | 0.23 | 0.33 | $3.70 \cdot 10^{-2}$ |
| **JUNB** | 0.13 | 0.08 | 0.54 | $4.82 \cdot 10^{-2}$ |

**Table S19.** Significantly differentially expressed DAM markers in the Mathys microglia for high AD association ($AD^{high}$) vs. low AD association ($AD^{low}$) microglia cells.

| | $AD^{high}$ expressed proportion | $AD^{low}$ expressed proportion | FC | t-test p-value |
|---|---|---|---|---|
| **CD74** | 0.55 | 0.31 | 1.53 | $1.08 \cdot 10^{-18}$ |
| **SPP1** | 0.53 | 0.22 | 1.62 | $5.71 \cdot 10^{-12}$ |
| **CTSB** | 0.36 | 0.25 | 0.78 | $6.89 \cdot 10^{-7}$ |
| **APOE** | 0.47 | 0.33 | 0.86 | $7.54 \cdot 10^{-7}$ |
| **TREM2** | 0.13 | 0.06 | 1.41 | $1.33 \cdot 10^{-4}$ |
| **RPLP2** | 0.23 | 0.15 | 0.85 | $2.07 \cdot 10^{-4}$ |
| **RPLP1** | 0.24 | 0.16 | 0.88 | $2.16 \cdot 10^{-4}$ |
| **HIF1A** | 0.26 | 0.19 | 0.66 | $8.37 \cdot 10^{-4}$ |
| **RPS19** | 0.39 | 0.34 | 0.44 | $8.57 \cdot 10^{-4}$ |
| **RPS20** | 0.21 | 0.15 | 0.74 | $9.82 \cdot 10^{-4}$ |
| **FTH1** | 0.34 | 0.24 | 0.67 | $2.59 \cdot 10^{-3}$ |
| **RPL13** | 0.34 | 0.26 | 0.45 | $3.76 \cdot 10^{-3}$ |

| | | | | |
|---|---|---|---|---|
| **RPS27A** | 0.21 | 0.13 | 0.66 | $4.22 \bullet 10^{-3}$ |
| **PABPC1** | 0.22 | 0.16 | 0.50 | $4.59 \bullet 10^{-3}$ |
| **RPS2** | 0.19 | 0.12 | 0.66 | $4.99 \bullet 10^{-3}$ |
| **NPC2** | 0.12 | 0.07 | 0.85 | $5.33 \bullet 10^{-3}$ |
| **RPL23** | 0.11 | 0.07 | 0.93 | $6.86 \bullet 10^{-3}$ |
| **RPL35** | 0.16 | 0.10 | 0.66 | $7.53 \bullet 10^{-3}$ |
| **RPS18** | 0.10 | 0.07 | 0.80 | $1.23 \bullet 10^{-2}$ |
| **RPL11** | 0.15 | 0.11 | 0.67 | $1.26 \bullet 10^{-2}$ |
| **TLR2** | 0.12 | 0.08 | 0.69 | $1.30 \bullet 10^{-2}$ |
| **RPS15** | 0.21 | 0.17 | 0.47 | $1.88 \bullet 10^{-2}$ |
| **RPL32** | 0.15 | 0.10 | 0.59 | $1.97 \bullet 10^{-2}$ |
| **RPL13A** | 0.24 | 0.20 | 0.42 | $1.97 \bullet 10^{-2}$ |
| **GNAS** | 0.21 | 0.17 | 0.37 | $2.76 \bullet 10^{-2}$ |
| **CLEC7A** | 0.12 | 0.07 | 0.62 | $3.14 \bullet 10^{-2}$ |
| **CSF2RA** | 0.30 | 0.26 | 0.28 | $3.60 \bullet 10^{-2}$ |
| **RPS24** | 0.16 | 0.12 | 0.46 | $4.80 \bullet 10^{-2}$ |
| **RPS14** | 0.15 | 0.11 | 0.45 | $4.89 \bullet 10^{-2}$ |

**Table S20.** Consistency of subtype clusters between multi-patient clustering (**Fig. 4A**) and single patient (**Fig. S2A-D**) clustering experiments. Rand, Fowlkes and Mallows's index (FM), and Jaccard index (JI) were used to measure the subtype cluster consistency.

| | **Rand** | **FM** | **JI** |
|---|---|---|---|
| **Patient 2** | 0.7059083 | 0.7446245 | 0.5872991 |
| **Patient 3** | 0.6362535 | 0.7976550 | 0.6362535 |
| **Patient 4** | 0.6064240 | 0.6417339 | 0.4611969 |
| **Patient 5** | 0.6683671 | 0.8059836 | 0.6637983 |

**Table S21.** Prediction performance metrics for IUSM myeloma single cells including ROC-AUC and PR-AUC.

| | **ROC-AUC** | **PR-AUC** |
|---|---|---|
| **Subtype 1** | 0.90 | 0.98 |
| **Subtype 2 (*PHF19$^{high}$*)** | 0.98 | 0.91 |
| **Subtype 3** | 0.90 | 0.62 |
| **Subtype 4** | 0.91 | 0.44 |
| **Subtype 5** | 0.98 | 0.84 |

**Table S22.** Spearman correlation coefficients between AUC for each of the patient output labels in patients and 100 randomly selected sets of hyperparameters. Significance values: • (0.1), * (0.05), ** (0.01), *** (0.001).

| | **Mesenchymal** | **Classical** | **Proneural** | **Neural** |
|---|---|---|---|---|
| **Training steps** | 0.20* | 0.11 | 0.19 | 0.17• |
| **Single cell batch size** | 0.03 | 0.07 | -0.02 | 0.01 |
| **Patient batch size** | 0.17• | 0.09 | 0.17• | 0.08 |
| **Hidden layer nodes** | 0.31** | 0.27** | 0.19• | 0.33*** |

| | | | | |
|---|---|---|---|---|
| **Drop-out retention rate** | 0.28** | 0.38*** | 0.42*** | 0.40*** |
| **Patient loss weight ($\lambda_1$)** | 0.03 | -0.08 | 0.03 | -0.04 |
| **MMD loss weight ($\lambda_2$)** | -0.18• | -0.12 | -0.10 | -0.05 |
| **L2 regularization weight ($\lambda_3$)** | -0.09 | -0.12 | -0.19• | -0.06 |

**Table S23.** Spearman correlation coefficients between correct label rank (MGH26: Proneural, MGH28: Mesenchymal, MGH29: Mesenchymal, MGH30: Classical) in single cells and 100 randomly selected sets of hyperparameters. Since there are 4 labels the best rank is 4, which indicates the correct label was assigned across the cells in that tumor. Significance values: • (0.1), * (0.05), ** (0.01), *** (0.001).

| | **MGH26** | **MGH28** | **MGH29** | **MGH30** |
|---|---|---|---|---|
| **Training steps** | -0.07 | -0.10 | -0.25* | -0.07 |
| **Single cell batch size** | -0.06 | 0.01 | 0.06 | 0.07 |
| **Patient batch size** | -0.06 | -0.22* | 0.01 | 0.06 |
| **Hidden layer nodes** | 0.10 | 0.27** | -0.08 | 0.06 |
| **Drop-out retention rate** | -0.07 | -0.06 | -0.34*** | 0.15 |
| **Patient loss weight ($\lambda_1$)** | 0.15 | -0.02 | 0.24* | 0.10 |
| **MMD loss weight ($\lambda_2$)** | 0.11 | -0.10 | 0.28* | 0.07 |
| **L2 regularization weight ($\lambda_3$)** | 0.18• | -0.09 | 0.01 | 0.03 |

**Table S24.** Prediction performance metrics for simulated patients including ROC-AUC and PR-AUC comparing *DEGAS* models with and without regularization.

| | **Simulation 1** | | **Simulation 2** | | **Simulation 3** | |
|---|---|---|---|---|---|---|
| | **ROC-AUC** | **PR-AUC** | **ROC-AUC** | **PR-AUC** | **ROC-AUC** | **PR-AUC** |
| **Disease status regularized** | 0.95 | 0.96 | 0.99 | 0.97 | 0.97 | 0.98 |
| **Disease status unregularized** | 0.94 | 0.96 | 0.99 | 0.97 | 0.94 | 0.93 |

**Table S25.** Prediction performance metrics for simulated cells including ROC-AUC and PR-AUC comparing *DEGAS* models with and without regularization.

| | **Simulation 1** | | **Simulation 2** | | **Simulation 3** | |
|---|---|---|---|---|---|---|
| | **ROC-AUC** | **PR-AUC** | **ROC-AUC** | **PR-AUC** | **ROC-AUC** | **PR-AUC** |
| **Cell type 1 regularized** | 1.00 | 1.00 | 1.00 | 1.00 | 1.00 | 1.00 |
| **Cell type 1 unregularized** | 0.95 | 0.93 | 1.00 | 1.00 | 0.90 | 0.75 |
| **Cell type 2 regularized** | 1.00 | 1.00 | 1.00 | 1.00 | 1.00 | 1.00 |
| **Cell type 2 unregularized** | 0.98 | 0.93 | 1.00 | 1.00 | 0.98 | 0.89 |
| **Cell type 3 regularized** | 1.00 | 1.00 | 1.00 | 1.00 | 1.00 | 1.00 |
| **Cell type 3 unregularized** | 0.93 | 0.86 | 1.00 | 1.00 | 0.91 | 0.79 |
| **Cell type 4 regularized** | 1.00 | 1.00 | 1.00 | 1.00 | 1.00 | 1.00 |

| | | | | | | |
|---|---|---|---|---|---|---|
| **Cell type 4 unregularized** | 0.95 | 0.93 | 1.00 | 1.00 | 0.89 | 0.78 |
